# Behavioural and neural signatures of perceptual evidence accumulation are modulated by pupil-linked arousal

**DOI:** 10.1101/433060

**Authors:** Jochem van Kempen, Gerard M. Loughnane, Daniel P. Newman, Simon P. Kelly, Alexander Thiele, Redmond G O’Connell, Mark A. Bellgrove

## Abstract

The timing and accuracy of perceptual decision making is exquisitely sensitive to fluctuations in arousal. Although extensive research has highlighted the role of neural evidence accumulation in forming decisions, our understanding of how arousal impacts these processes remains limited. Here we isolated electrophysiological signatures of evidence accumulation alongside signals reflecting target selection, attentional engagement and motor output and examined their modulation as a function of both tonic and phasic arousal, indexed by baseline and task-evoked pupil diameter, respectively. For both pupillometric measures, the relationship with reaction time was best described by a second-order, U-shaped, polynomial. Additionally, the two pupil measures were predictive of a unique set of EEG signatures that together represent multiple information processing steps of perceptual decision-making, including evidence accumulation. Finally, we found that behavioural variability associated with fluctuations in both tonic and phasic arousal was largely mediated by variability in evidence accumulation.

## Introduction

The speed and accuracy with which humans, as well as non-human animals, respond to a stimulus depends not only on the characteristics of the stimulus, but also on the cognitive state of the subject. When drowsy, a subject will respond more slowly to the same stimulus compared to when she is attentive and alert. Central arousal also fluctuates across a smaller range during quiet wakefulness, when the subject is neither drowsy or inattentive, nor overly excited or distractible. Although these trial-to-trial fluctuations can impact on behavioural performance during decision-making tasks (Aston-Jones and Cohen, 2005), it is largely unknown how arousal modulates the underlying processes that support decision formation. Perceptual decision-making depends on multiple neural processing stages that represent and select sensory information, those that process and accumulate sensory evidence, and those that prepare and execute motor commands. Variability in central arousal could affect any one or potentially all of these processing stages, which in turn could influence behavioural performance.

The neuromodulatory systems that control central arousal state, such as the noradrenergic (NA) locus coeruleus (LC) and the cholinergic basal forebrain (BF), have also been suggested to drive fluctuations in endogenous activity linked to changes in cortical (de)synchronization, i.e. cortical state (Harris and Thiele, 2011; Lee and Dan, 2012), and are linked to cognitive functions such as attention (Thiele and Bellgrove, 2018), both known to affect information processing and behavioural performance. These modulatory systems have both tonic and phasic firing patterns that are recruited on different timescales and support different functional roles (Aston-Jones and Cohen, 2005; Dayan and Yu, 2006; Parikh et al., 2007; Parikh and Sarter, 2008; Sarter et al., 2016). Tonic changes in neuromodulator activity occur over longer timescales that can span multiple trials, whereas fast (task-evoked) recruitment through phasic activation occurs on short enough timescales to influence neural activity and behavioural decisions within the same trial (Aston-Jones and Cohen, 2005; Bouret and Sara, 2005; Dayan and Yu, 2006; Parikh et al., 2007).

Pupil diameter correlates strongly with a variety of measurements of cortical state and behavioural arousal (Eldar et al., 2013; Reimer et al., 2014; McGinley et al., 2015b, 2015a; Vinck et al., 2015; Engel et al., 2016), and can thus be considered a reliable proxy of central arousal state. Indeed, there is a strong correlation between pupil size and activity in various neuromodulatory centres that control arousal (Aston-Jones and Cohen, 2005; Gilzenrat et al., 2010; Murphy et al., 2014a; Varazzani et al., 2015; Joshi et al., 2016; Reimer et al., 2016; de Gee et al., 2017). Both baseline pupil diameter, reflecting tonic activity levels in neuromodulatory centres (tonic arousal), and task-evoked pupil diameter changes (phasic arousal), have been related to specific neural processing stages of perceptual decision making. Baseline pupil diameter correlates with sensory sensitivity (McGinley et al., 2015a, 2015b) and is predictive of behavioural performance during elementary detection tasks (Murphy et al., 2011; McGinley et al., 2015a). Pupil diameter also changes phasically in the course of a single decision (Beatty, 1982a; de Gee et al., 2014, 2017; Lempert et al., 2015; Murphy et al., 2016; Urai et al., 2017), and has been related to specific elements of the decision making process, such as decision bias (de Gee et al., 2014, 2017), uncertainty (Urai et al., 2017), and urgency (Murphy et al., 2016). This suggests that these neuromodulatory systems do not only dictate network states (through tonic activity changes), but that they are recruited throughout the decision making process (Cheadle et al., 2014; de Gee et al., 2014, 2017). Although both baseline pupil diameter and the phasic pupil response have been associated with specific aspects of decision-making, the relationship between pupil-linked arousal and the electrophysiological correlates of decision-making, and in particular evidence accumulation, are largely unknown.

Recently developed behavioural paradigms have made it possible to non-invasively study the individual electroencephalographic (EEG) signatures of perceptual decision-making described above (O’Connell et al., 2012; Kelly and O’Connell, 2013; Loughnane et al., 2016, 2018; Newman et al., 2017). In these paradigms, participants are required to continuously monitor (multiple) stimuli for subtle changes in a feature. Because stimuli are presented continuously, target onset times (and locations) are unpredictable, and sudden stimulus onsets are absent, eliminating sensory evoked deflections in the EEG traces. These characteristics allow for the investigation of the gradual development of build-to-threshold decision variables as well as signals that code for the selection of relevant information from multiple competing stimuli, a critical feature of visuospatial attentional orienting that impact evidence accumulation processes (Loughnane et al., 2016).

Here, we asked how arousal influences EEG signals that relate to each of the separate processing stages described above. Specifically, we tested the effects of pupil-linked arousal on pre-target preparatory parieto-occipital α-band activity, associated with fluctuations in the allocation of attentional resources (Kelly and O’Connell, 2013); early target selection signals measured over contra- and ipsilateral occipital cortex, the N2c and N2i (Loughnane et al., 2016); perceptual evidence accumulation signals measured as the centroparietal positivity (CPP), which is a build-to-threshold decision variable demonstrated to scale with the strength of sensory evidence and predictive of reaction time (RT) (O’Connell et al., 2012; Kelly and O’Connell, 2013); and motor-preparation signals measured via contralateral β-band activity (Donner et al., 2009; O’Connell et al., 2012). Of these signals, we extracted specific characteristics such as the latency, build-up rate and amplitude, and tested whether these were affected by pupil-linked arousal. Additionally, because the variance and response reliability of the membrane potential of sensory neurons varies with pupil diameter (Reimer et al., 2014; McGinley et al., 2015a), we also investigated whether arousal affected the inter-trial phase coherence (ITPC), a measure of across trial consistency in the EEG signal, of the N2 and the CPP.

We found that both baseline pupil diameter as well as the pupil response were predictive of behavioural performance, and that this relationship was best described by a U-shaped, second-order polynomial, model fit. Furthermore, we found that both tonic and phasic arousal bore a predictive relationship with the neural signals coding for baseline attentional engagement, early target selection, evidence accumulation as well as the preparatory motor response. Although neural activity representing all these stages varied with changes in arousal, unique variability in task performance due to tonic arousal (baseline pupil diameter) could only be explained by the amplitude of target selection signals and the consistency of the build-up rate of the CPP, reflecting evidence accumulation. In contrast, variability due to phasic arousal (pupil response) was explained by pre-target α-band activity as well as the build-up rate and consistency of the CPP.

## Results

80 subjects performed a continuous version of the random dot motion task in which they were asked to report temporally and spatially unpredictable periods of coherent motion within either of two streams of random motion (Figure 1A). We investigated whether the trial-to-trial fluctuations in behavioural performance and EEG signatures of perceptual decision making could, in part, be explained by trial-to-trial differences in the size of the baseline pupil diameter (reflecting tonic arousal) and the post-target pupil response (reflecting phasic arousal). We quantified this relationship by allocating data into 5 bins based on the size of either the baseline pupil diameter or the phasic pupil diameter response (Figure 1B & Figure 1D). We then used sequential multilevel model analyses and maximum likelihood ratio tests to test for fixed effects of pupil bin. We determined whether a linear fit was better than a constant fit and subsequently whether the fit of a second-order polynomial (e.g, U-shaped relationship), indicating a non-monotonic relationship between pupil diameter and behaviour/EEG, was superior to a linear fit.

**Figure 1.**
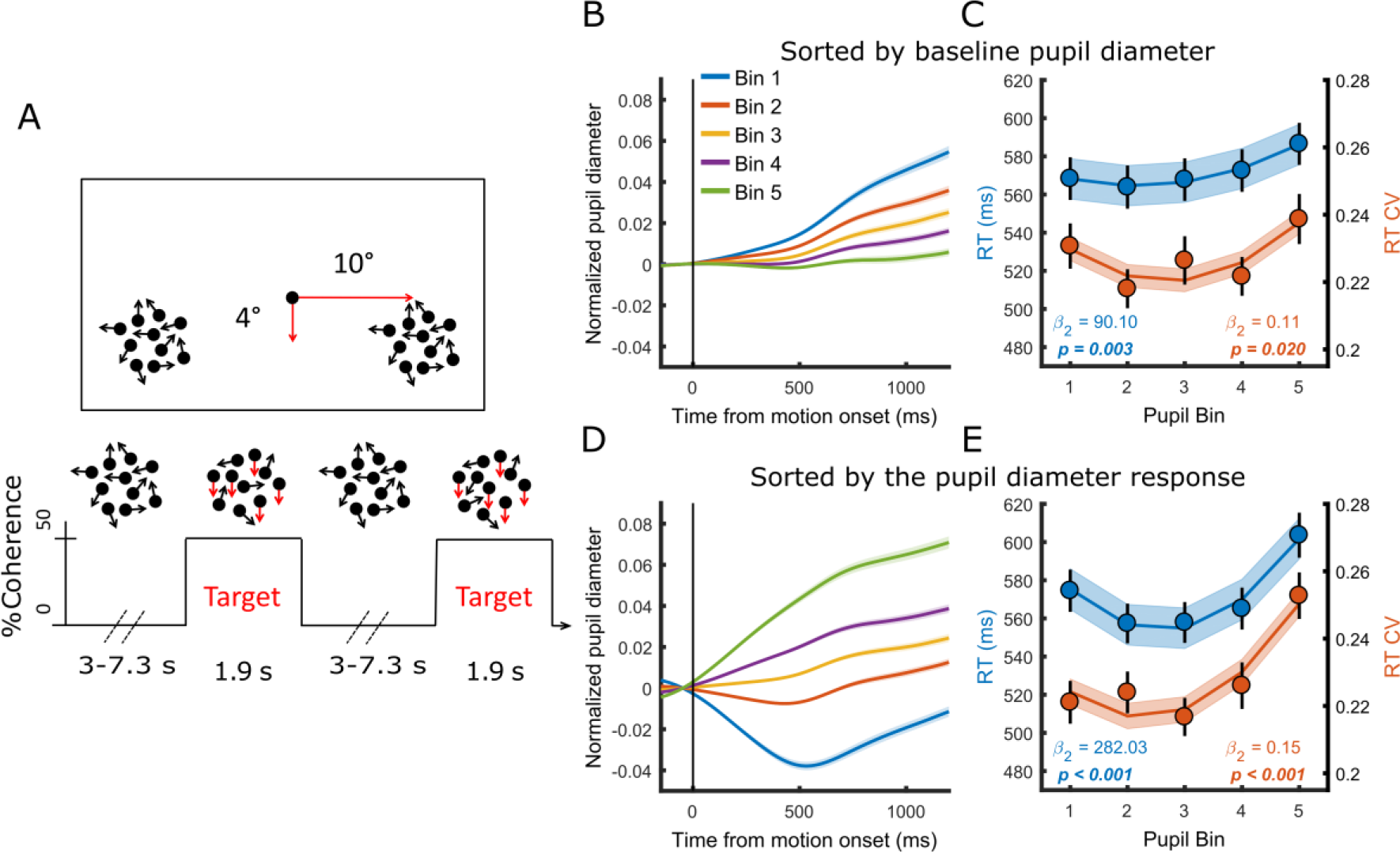
(A) Paradigm. Subjects fixated on a central dot while monitoring two peripheral patches of continuously presented randomly moving dots. At pseudorandom times an intermittent period of coherent downward motion (50%) occurred in either the left or the right hemifield. A speeded right handed button press was required upon detection of coherent motion. (B-C) Pupil diameter time course and task performance sorted by baseline pupil diameter. (B) Pupil time-course for the five bins. (C) Behavioural performance for the five bins. Markers indicate mean reaction times (RT, blue, left y-axis) and reaction time coefficient of variation (RTcv, red, right y-axis), lines and shading indicate significant quadratic fits. (D-E) Same conventions as B-C, but sorted by the pupil diameter response. Error bars and shaded regions denote ±1 standard error of the mean (SEM). Stats, linear mixed effects model analyses (Statistical analyses).

### Both tonic and phasic arousal are predictive of task performance in a U-shaped manner

We first investigated the relationship between trial-by-trial pupil dynamics and behavioural performance. As stimuli were presented well above perceptual threshold, our subjects performed at ceiling (Newman et al., 2017). We therefore focused on RT and the RT coefficient of variation (RTcv), a measure of performance variability calculated by dividing the standard deviation in RT by the mean (Bellgrove et al., 2004), rather than accuracy. We found that both measures of behavioural performance displayed a non-monotonic, U-shaped, relationship with both baseline pupil diameter (RT χ^2^_(1)_ = 8.98, p = 0.003; RTcv χ^2^_(1)_ = 5.36, p = 0.020) and the pupil diameter response (RT χ^2^_(1)_ = 116.65, p < 0.001; RTcv χ^2^_(1)_ = 12.36, p < 0.001). Responses were fastest and least variable for intermediate pupil bins (Figure 1C & Figure 1E). We repeated this analysis in single-trial, non-binned data, in which we additionally controlled for time-on-task effects, confirming that these effects were not dependent on the binning procedure (Supplementary information). Additionally, we noticed that when we band-pass filtered the pupil diameter, rather than low-pass filtered, the relationship between baseline pupil diameter and task performance was not significant, whereas this did not affect the relationship between the pupil response and task performance (Supplementary figure 1). This suggests that slow fluctuations in baseline pupil diameter (<0.01Hz) are driving the effect on task performance.

Having established a relationship between task performance and both tonic and phasic modes of central arousal state, we next focused on the relationship between these pupil dynamics and the neural signatures underpinning target detection on this perceptual decision making task (Loughnane et al., 2016; Newman et al., 2017).

### U-shaped relationship between phasic arousal and decision computation

During decision making, perceptual evidence has to be accumulated over time. This accumulation process has long been related to build-to-threshold activity in single neurons in parietal cortex (Gold and Shadlen, 2007; but see Latimer et al., 2015, 2016; Shadlen et al., 2016). The centro-parietal positivity (CPP) measured from scalp EEG exhibits many of these same properties, including a representation of the accumulation of sensory evidence towards a decision bound (O’Connell et al., 2012, 2018; Kelly and O’Connell, 2013). Here we tested the relationship between the pupil diameter response and the onset, build-up rate, amplitude and consistency (ITPC) of the CPP (Figure 2). We found that the onset latency of the evidence accumulation process, defined as the first time point that showed a significant difference from zero for 15 consecutive time points, displayed a quadratic relationship with the size of the pupil response (χ^2^_(1)_ = 7.53, p = 0.006), such that the fastest onsets were found for intermediate pupil response bins and slower onsets for the extreme bins (Figure 2A). Likewise, the slope of the CPP, reflecting the build-up rate of evidence accumulation, also displayed a non-monotonic, inverted-U shaped, relationship with the pupil response (χ^2^_(1)_ = 7.81, p = 0.005). The amplitude of the CPP, representing the threshold of the accumulation process, did not vary with the pupil diameter response (p = 0.24). We thus found a direct relationship between phasic arousal and the onset and build-up rate of evidence accumulation. Moreover, the non-monotonic relationship with the neural parameters of the CPP closely resembled the relationship between the pupil response and behavioural performance (Figure 1E). Because the membrane potential of sensory neurons shows the least variance and highest response reliability at intermediate baseline pupil diameter (McGinley et al., 2015a), we additionally investigated the ITPC, a measure of across trial consistency, of the CPP. We computed ITPC with a single-taper spectral analysis in a 512 ms sliding window computed at 50 ms intervals, with a frequency resolution of 1.95 Hz (Materials and Methods). Based on the stimulus-locked grand average time-frequency spectrum, we selected a time (300-550 ms) and frequency window (<4 Hz) for further statistical analyses (Figure 2C). We found a quadratic (inverted U-shape) relationship between pupil diameter response and the consistency of the CPP signal (χ^2^_(1)_ = 30.42, p < 0.001), indicating that the CPP signal is less variable for intermediate pupil response bins (Figure 2D). Together, these results confirm the hypothesized relationship between the pupil diameter response and electrophysiological correlates of evidence accumulation. Next, we asked whether other stages of information processing underpinning perceptual decision making also varied with the pupil response.

**Figure 2.**
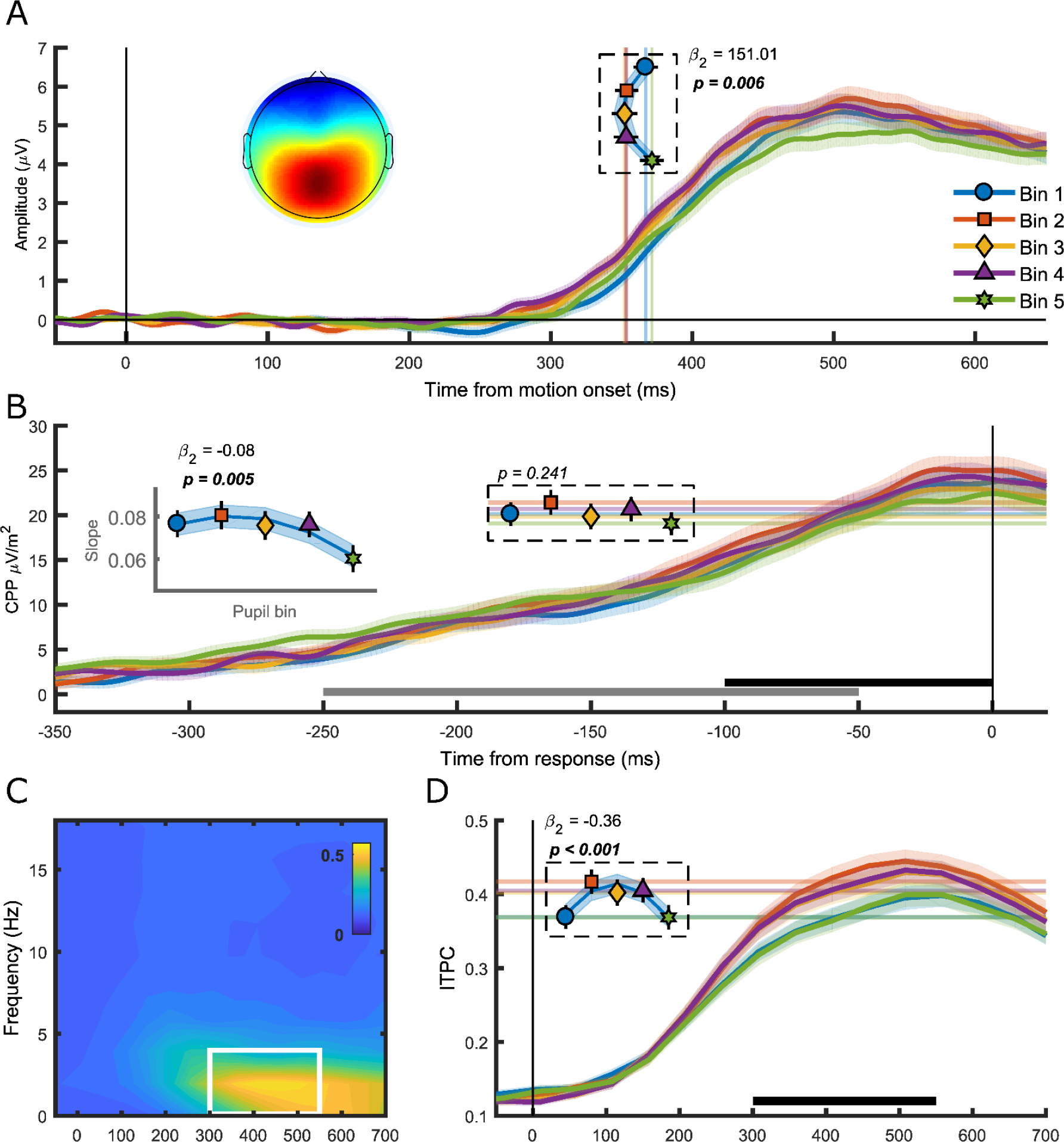
The centro-parietal positivity (CPP) in relation to phasic arousal. (A) The stimulus-locked CPP time-course shows faster onset times for intermediate pupil response bins. The inset shows the scalp topography of the CPP. Vertical lines and markers indicate the onset latencies per bin. (B) The response-locked CSD-transformed CPP time-course. Horizontal lines and symbols indicate the CPP amplitude, and the inset displays the build-up rate of the CPP across pupil response bins. The black bar represents the time window used for the calculation of the CPP amplitude and the grey bar the time window used for the calculation of the build-up rate. (C) Grand average inter-trial phase coherence (ITPC) per time-frequency point for the CPP. White box represents the time-frequency window selected for statistical analyses. (D) ITPC per pupil bin over time for frequencies below 4 Hz. The black bar indicates the time window used for further analysis. Horizontal lines and symbols indicate the averaged ITPC in the time-frequency window indicated by the white box in panel C. Lines and shading indicate significant quadratic fits to the data. Error bars and shaded regions denote ±1 SEM. Stats, linear mixed effects model analyses (Statistical analyses).

### The phasic pupil response relates monotonically to spectral measures of baseline attentional engagement and displays a U-shaped relationship with motor output

We next investigated pre-target preparatory α-band power (8-13 Hz), a sensitive index of attentional deployment that has been shown to vary with behavioural performance. Specifically, previous studies have found higher pre-target α-band power preceding trials with longer RT, and that fluctuations in α-power may reflect an attentional influence on variability in task performance (Ergenoglu et al., 2004; van Dijk et al., 2008; O’Connell et al., 2009; Kelly and O’Connell, 2013). We first verified the relationship between α-band power and behavioural performance by binning the data into 5 bins according to α-band power and performing the same sequential regression analysis as described above (Figure 3A). We replicated previous findings (Kelly and O’Connell, 2013) and found an approximately linear relationship between α-band power and RT (χ^2^_(1)_ = 23.31, p < 0.001) but not RTcv (p = 0.26). In line with previous research (Hong et al., 2014), pupil diameter response was inversely related to α-band power (Figure 3B), displaying an approximately linear relationship (χ^2^_(1)_ = 47.19, p < 0.001), suggesting that pre-target attentional engagement is related to phasic arousal.

We next focused on response-related motor activity in the form of left hemispheric β-power (LHB). LHB decreases before a button press and has been shown to reflect the motor-output stage of perceptual decision making, but also to trace decision formation, reflecting the build-up of choice selective activity (Donner et al., 2009). Here we investigated the LHB amplitude and build-up rate preceding response (Figure 3C). We found that both LHB amplitude (χ^2^_(1)_ = 4.18, p = 0.041) and LHB slope (χ^2^_(1)_ = 3.94, p = 0.047) displayed a non-monotonic relationship with pupil response, suggesting that phasic arousal influences the build-up rate of choice-related activity over motor cortex. The build-up rate results accord with those for the CPP, as for both measures the slope declined with a larger pupil diameter response (note that LHB decreases during decision formation, i.e. has a negative slope).

### Only ipsilateral target selection signals correlate with the phasic pupil response

Next we investigated the N2 (Figure 3D-F), a stimulus-locked early target selection signal that has been shown to predict behavioural performance and modulate the onset and build-up rate of the CPP (Loughnane et al., 2016). Because of the spatial nature of the task, we analysed the negative deflection over both the contra-(N2c) and ipsi-lateral (N2i) hemisphere, relative to the target location. The pupil response was not predictive of any aspect of the N2c. Specifically, phasic arousal was not predictive of N2c latency (p = 0.66) or amplitude (p = 0.39), nor did we find any relationship between the pupil response and the N2c ITPC (p = 0.57). Although the pupil response was not predictive of N2i latency (p = 0.53) or ITPC (0.69), it was predictive of N2i amplitude (χ^2^_(1)_ = 6.94, p = 0.008). Previously, we showed that the N2c, rather than N2i, correlated with RT and modulated CPP (Loughnane et al., 2016). It is therefore interesting that N2i, rather than N2c varied with the pupil response. Below, we will discuss whether this effect could (partially) explain the relationship between the pupil response and task performance.

**Figure 3.**
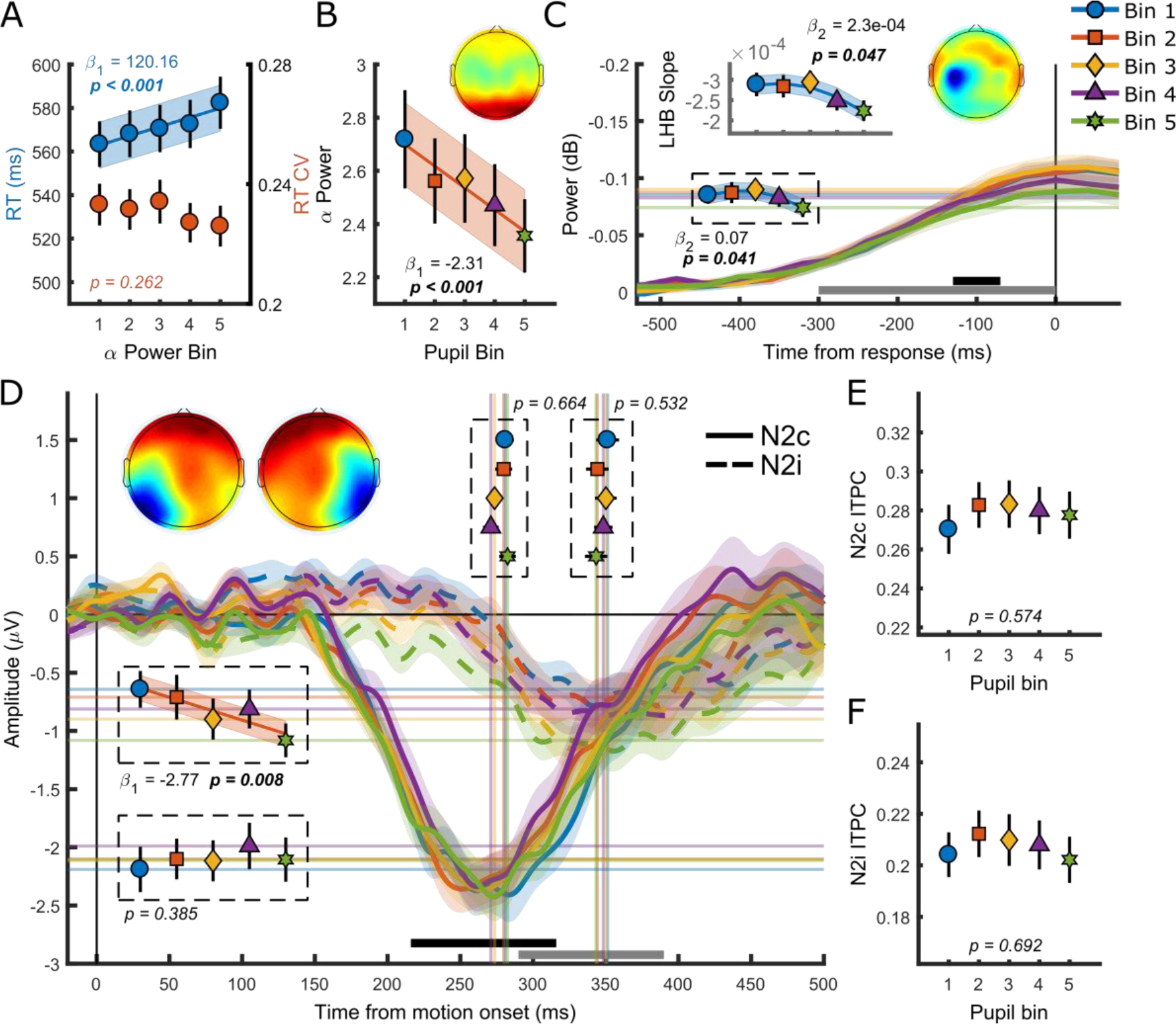
(A) RT and RTcv in relation to pre-target α power. (B) Pre-target α power in relation to the pupil response. (C) response-related left hemispheric β power (LHB) per pupil bin. Horizontal lines and marks indicate the average LHB in the time window indicated by the black bar. Inset shows LHB build-up rate, as determined by fitting a straight line through the LHB in the time window indicated by the grey bar. Note the reverse y-axis direction. (D) The stimulus-locked N2c (solid lines) and N2i (dashed lines) time-course binned by the pupil response. Vertical lines and markers show the peak latencies. Horizontal lines and markers show the average N2 amplitude during the time period indicated by the black (N2c) and grey (N2i) bars. (E-F) N2c (E) and N2i (F) ITPC per pupil bin over the time and frequency window determined based on the grand average ITPC (Supplementary figure 2). Insets show the scalp topography of each neural signal. Lines and shading indicate significant fits to the data. Linear fits are displayed when a first-order polynomial fit was superior to a constant fit, and quadratic fits are displayed where second-order fits were superior to a first-order fit. Error bars and shaded regions denote ±1 SEM. Stats, linear mixed effects model analyses (Statistical analyses).

### Variation in spatial attention influences task performance, but cannot explain the U-shaped relationship between pupil response and RT

Having established a relationship between the size of the pupil response and both task performance and EEG signatures of perceptual decision making, we investigated whether the U-shaped relationship with behavioural performance could be explained by factors other than phasic arousal. Alongside activity in neuromodulatory centres, pupil diameter also correlates with activity in the intermediate layers of the superior colliculus (SCi) (Wang et al., 2012; Joshi et al., 2016; de Gee et al., 2017). The SCi, besides preparing and executing eye movements, is involved in directing covert attention (Kustov and Lee Robinson, 1996; Ignashchenkova et al., 2004; Muller et al., 2005; Lovejoy and Krauzlis, 2010), and provides an essential contribution to the selection of stimuli from amongst competing distractors (McPeek and Keller, 2002, 2004; reviewed in Mysore and Knudsen, 2011). Therefore, given our use of multiple simultaneously presented competing stimuli, variations in spatial attention could potentially explain variability in behavioural performance and pupil diameter responses. Indeed, previous research has reported an association between poorer behavioural performance and large pupil diameter responses when there was a requirement to monitor multiple stimuli simultaneously (Kristjansson et al., 2009).

To test this possibility, we further investigated the relationship between pupil responses and the ipsilateral N2 target selection signal (Figure 3D). If on trials with lower behavioural performance attention was focused on the distractor stimulus, then early target selection signals contralateral to the distractor stimulus (i.e. ipsilateral to the target stimulus) might differ compared to trials with relatively better performance. Additionally, these differences might be present throughout the trial, before the N2i is expected to reveal differences between target and non-target stimuli (Loughnane et al., 2016). We therefore conducted a sliding window linear mixed effect model analysis predicting N2i amplitude for each 100 ms window, in 10 ms increments, from −20 before to 500 ms after target onset (Figure 4A). This analysis revealed that the pupil diameter was predictive of N2i amplitude from as early as 70 ms after target onset, much earlier than the previously reported target selection onset of 308 ms (Loughnane et al., 2016), and therefore unlikely to reflect target processing. Rather, large pupil responses and a large N2 amplitude could reflect a bias in attention or expectation of the target location. Although large pupil responses have previously been related to a reduction in decision bias (de Gee et al., 2014, 2017), these studies did not investigate decision bias in the context of spatially unpredictable target locations. As the size of the N2i could indicate the need to shift attention, we next tested whether the size of the N2i amplitude could predict the size of the pupil diameter response. In line with the results described above, where trials with larger pupil responses displayed larger N2i amplitudes, N2i amplitude displayed an inverse relationship (note that the first bin contains the largest N2i responses) with the pupil diameter response (χ^2^_(1)_ = 6.91, p = 0.009), with larger pupil responses for larger N2i amplitudes (Figure 4B). This suggests that attentional shifts, possibly through recruitment of the SCi, could lead to larger pupil responses and lower behavioural performance.

**Figure 4.**
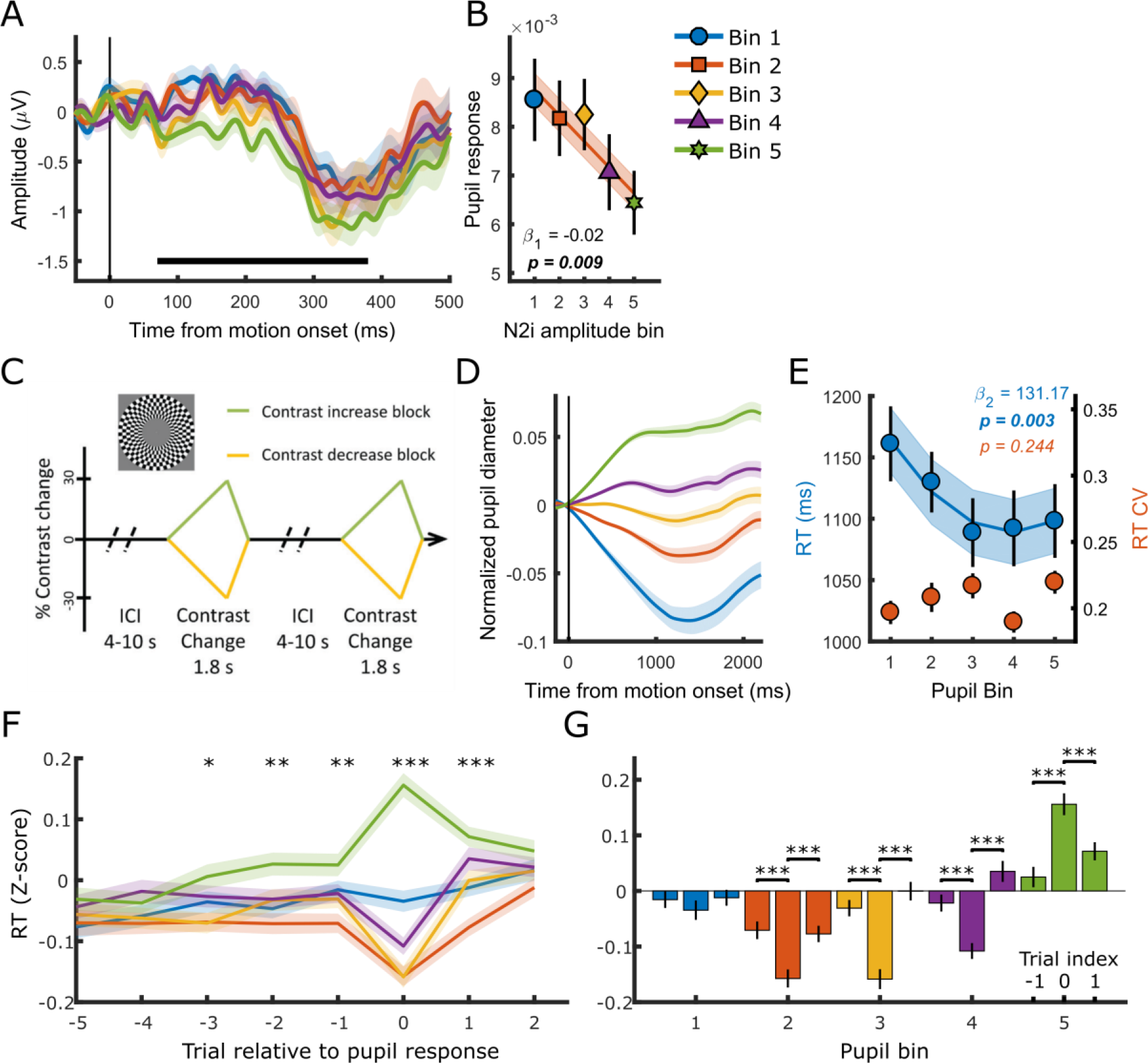
(A) The stimulus-locked N2i time-course binned by the pupil response. The time period indicated by the black bar displays the times where there was a significant, approximately linear, relationship between the pupil response and N2i amplitude. Stats, sliding window linear mixed effect model analysis (main text), FDR corrected. (B) The size of the pupil response, sorted by N2i amplitude. (C-E) Data from a different dataset using a contrast change detection paradigm where subjects monitored a single central target stimulus (Loughnane et al., 2018). (C) Task paradigm. Participants monitored a single central checkerboard stimulus for gradual contrast changes. (D) Pupil time-courses for the 5 bins. (E) Relationship between behavioural performance and the pupil diameter response. Markers indicate mean reaction times (RT, blue, left y-axis) and reaction time coefficient of variation (RTcv, red, right y-axis). (F) Z-scored log transformed RT across subsequent trials, sorted by pupil diameter on trial 0. Significance stars indicate when RT for the 5^th^ pupil bin was significantly higher than the average RT across the first 4 bins. (G) RT per pupil bin for trial index −1, 0 and 1 (panel F). Lines and shading indicate significant fits to the data. Linear fits are displayed when a first-order polynomial fit was superior to a constant fit, and quadratic fits are displayed where second-order fits were superior to a first-order fit. Error bars and shaded regions denote ±1 SEM. Stats, linear mixed effects model analyses (panel A-E, Statistical analyses), and paired sample t-tests, FDR corrected (panel F-G).

To further investigate whether these effects could explain (part of) the current results, we analysed the relationship between the pupil diameter response and task performance from a different dataset (Loughnane et al., 2018) in which participants (n = 17) monitored a single, centrally positioned, flickering checkerboard annulus for a gradual change in contrast (Figure 4C). Pupil diameter on this non-spatial task also displayed across-trial variability (Figure 4D), which predicted RT in a non-monotonic fashion (χ^2^_(1)_ = 8.85, p = 0.003). RTcv did not scale with the pupil diameter response (Figure 4E) (p = 0.24). We furthermore confirmed that the non-monotonic relationship between the pupil response and RT was not dependent on the binning procedure or time-on-task effects by repeating this analysis on single-trial data in which we controlled for these factors (Supplementary Table 1). Thus, the U-shaped relationship between the pupil diameter response and behaviour (RT) cannot be attributed to attentional shifts away from a distractor stimulus and may be a more general phenomenon during protracted perceptual decision-making.

### Large pupil responses may indicate a performance monitoring compensatory mechanism

For both NA and acetylcholine (ACh) it has been found that phasic activity is task dependent and generally larger on trials with good performance (Aston-Jones et al., 1994, 1997, Rajkowski et al., 1994, 2004; Parikh et al., 2007; Gritton et al., 2016). It therefore seems counterintuitive that in this study large pupil responses, presumably reflecting transient activity in these modulatory systems amongst others, are associated with lower behavioural performance. One possible explanation of these results is that this transient reflects a NA and/or ACh-related compensatory mechanism (Murphy et al., 2011; Sarter et al., 2016). Although speculative, a transient in phasic arousal could, on the current task, reflect a correction from a state with low performance. Indeed, trials with maximum pupil dilations and low task performance have been found to be preceded by trials with progressively longer RT, and followed by better task performance (Murphy et al., 2011). We therefore tested whether trials with a large pupil diameter response were preceded/followed by trials with worse/better task performance. Figure 4F-G shows the RT for trials relative to the trial on which the pupil response was measured (trial 0). Trials with larger pupil responses (bin 5, green) were preceded by trials with slower than average RT (Figure 4F), and this effect was observed for up to 3 trials before trial 0. Although on the subsequent trial (trial 1) RT was still slower than average, RT was significantly faster compared to trial 0 (Figure 4G). Additionally, the trials for which the pupil response is largest are the only trials on which there is both a decrease in task performance (increase in RT) from the previous trial, and a subsequent improvement in performance on the next trial. The other bins displayed the exact opposite pattern, and none of them showed an improvement in task performance after trial 0. A phasic pupillary response could thus indicate a compensatory mechanism, signalling the need to adjust the neural circuitry to a state that facilitates better performance. As Murphy et al. (2011) concluded, large pupil responses may reflect phasic activations driven by higher cortical performance monitoring brain regions that serve to reengage participants in the task.

### The impact of phasic arousal on task performance is mainly mediated by the consistency in evidence accumulation

Regardless of the neural mechanism, we found that pupil-linked phasic arousal was predictive of specific neural signals at multiple information processing stages of perceptual decision making. To test which of these signals explained unique variability in behavioural performance across the 5 pupil response bins and subjects, the neural signals were added to a linear mixed effects model predicting either RT or RTcv with their order of entry determined hierarchically by their temporal order in the decision-making process. This allowed us to test whether each successive stage of neural processing would improve the fit of the model to the behavioural data, over and above the fit of the previous stage. Note that none of the predictors were highly correlated (r < 0.25), with the exception of CPP onset and CPP ITPC (r = 0.43), CPP build-up rate and CPP amplitude (r = −0.59), and LHB build-up rate and amplitude (r = −0.28). Compared to the baseline model predicting RT with pupil bin, the addition of pre-target α-power significantly improved the model fit (χ^2^_(1)_ = 10.63, p < 0.001). None of the measures of early target selection improved the fit of the model; neither N2c latency (χ^2^_(1)_ = 0.75, p = 0.39) or amplitude (χ^2^_(1)_ = 0.47, p = 0.49), nor N2i latency (χ^2^_(1)_ = 0.90, p = 0.34) or amplitude (χ^2^_(1)_ = 2.34, p = 0.13). We found that both the addition of CPP onset (χ^2^_(1)_ = 27.24, p < 0.001) as well as the build-up rate (χ^2^_(1)_ = 11.74, p < 0.001) significantly improved the model fit. Whereas the addition of CPP amplitude did not (χ^2^_(1)_ = 3.19, p = 0.07), the addition of CPP ITPC substantially improved the fit of the model (χ^2^_(1)_ = 40.60, p < 0.001). Although both LHB amplitude and build-up rate varied with phasic arousal, neither improved the fit of the model (LHB build-up rate χ^2^_(1)_ = 2.09, p = 0.15; amplitude χ^2^_(1)_ = 0.59, p = 0.44). Overall, this model suggested that pre-target α-power, CPP onset, build-up rate and ITPC exert partially independent influences on RT. Because some variables were highly correlated (e.g. CPP onset and ITPC) we used an algorithm for forward/backward stepwise model selection (Venables and Ripley, 2002) to test whether each neural signal indeed explained independent variability that is not explained by any of the other signals. This procedure eliminated CPP onset from the final model (F_(1)_ = 2.60, p = 0.11). Thus, only pre-target α-power, CPP build-up rate and CPP ITPC significantly improved the model fit for predicting RT. These three variables were forced into one linear mixed effects model predicting RT (Statistical analyses), and comparison to a baseline model revealed a good fit (χ^2^_(3)_ = 82.18, p < 0.001). The fixed effects of the model (the neural signals) explained 14.6% of the variability in RT (marginal *r*^2^) across the 5 pupil response bins, and together with the random effects (across subject variability) it explained 93.1% of the variability (conditional *r*^2^).

We performed the same hierarchical regression analysis to see which neural signals explained variability in RTcv. We summarised the results of this analysis in Supplementary Table 2, and report the most important results here. The hierarchical regression analysis revealed that both CPP onset and CPP ITPC improved the model fit, but eliminated CPP onset after the forward/backward model selection. Consequently, CPP ITPC was the only variable that exerted independent influence on RTcv. Comparison against a baseline model revealed a significant fit (χ^2^_(1)_ = 19.78, p < 0.001) that had a marginal *r*^2^ of 11.1% and a conditional *r*^2^ of 46.5%.

Table 1 shows the final parameter estimates for the neural signals that significantly predicted variability in RT or RTcv that is due to variability in phasic arousal. From this analysis we can conclude that CPP ITPC was the strongest predictor for RT and the only predictor for RTcv. These results provide novel insight into the mechanism by which the neuromodulators that control arousal can influence behaviour. The impact of these modulators on decision-making, previously suggested to be recruited throughout the decision-making process (Cheadle et al., 2014; de Gee et al., 2014, 2017), is thus mainly mediated by their effects on the consistency in evidence accumulation.

Next, we turn to tonic arousal and its relationship to these same EEG components of perceptual decision-making.

**Table 1.** Parameter estimates for the final linear mixed effect model of RT and RTcv binned by the pupil diameter response or baseline. The only parameters included are the neural signals that significantly improved the model fit.

|  | RT |  |  |  | RTcv |  |  |  |
| --- | --- | --- | --- | --- | --- | --- | --- | --- |
| | $\beta$ | $\beta$ SE | T | p | $\beta$ | $\beta$ SE | t | P |
| *Pupil response* |  |  |  |  |  |  |  |  |
| **pre-target α-power** | 0.18 | 0.050 | 3.70 | <0.001 |  |  |  |  |
| **CPP build-up rate** | -0.10 | 0.042 | -2.26 | 0.024 |  |  |  |  |
| **CPP ITPC** | -0.23 | 0.029 | -7.97 | <0.001 | -0.26 | 0.056 | -4.66 | <0.001 |
| *Baseline Pupil diameter* |  |  |  |  |  |  |  |  |
| **N2c amplitude** | 0.05 | 0.027 | 1.95 | 0.052* |  |  |  |  |
| **CPP ITPC** | -0.18 | 0.032 | -5.48 | <0.001 | -0.31 | 0.056 | -5.45 | <0.001 |

\* Although N2c amplitude fell short of the nominal statistical significance threshold, a robust regression analysis confirmed a significant positive relationship with RT (Supplementary Table 4).

### Baseline pupil diameter is inversely related to the consistency of evidence accumulation

Figure 5 illustrates the relationship between baseline pupil diameter and the CPP. Unlike the pupil response, baseline pupil diameter was not predictive of the onset (p = 0.17), build-up rate (p = 0.15), or amplitude of the CPP (p = 0.10). The only component that significantly scaled with baseline pupil diameter was the consistency of evidence accumulation, CPP ITPC (χ^2^_(1)_ = 9.34, p = 0.002). In line with previous research that revealed increased variability in the rate of evidence accumulation during periods with larger baseline pupil diameter (Murphy et al., 2014b), we found an inverse, approximately linear, relationship in which higher baseline pupil diameter displayed lower EEG signal consistency (Figure 5D). Thus, states of higher arousal are characterized by less consistency, i.e. more variability, in the accumulation of evidence. Additionally, these states (bin 4 and 5) also show slower RT and higher RTcv (Figure 1C), indicating that higher variability in the rate of evidence accumulation (due to higher tonic arousal) affects task performance.

### Baseline pupil diameter relates to spectral measures of baseline attention engagement and motor output as well as early target selection

We found a relationship between baseline pupil diameter and specific characteristics of multiple neural processing stages of perceptual decision-making. Specifically, as observed before (Hong et al., 2014), pre-target alpha power (Figure 6A) varied with baseline pupil diameter in a non-monotonic, inverted-U shaped, manner (χ^2^_(1)_ = 4.40, p = 0.036). This suggests that with higher tonic arousal, alpha activity is higher (or less desynchronised). Next, we tested whether baseline pupil diameter was predictive of EEG characteristics representing motor output (Figure 6B). We found an approximately linear relationship with LHB build-up rate (χ^2^_(1)_ = 11.1, p < 0.001), decreasing with larger baseline pupil diameter, but we did not find a relationship with LHB amplitude (p = 0.18).

**Figure 5.**
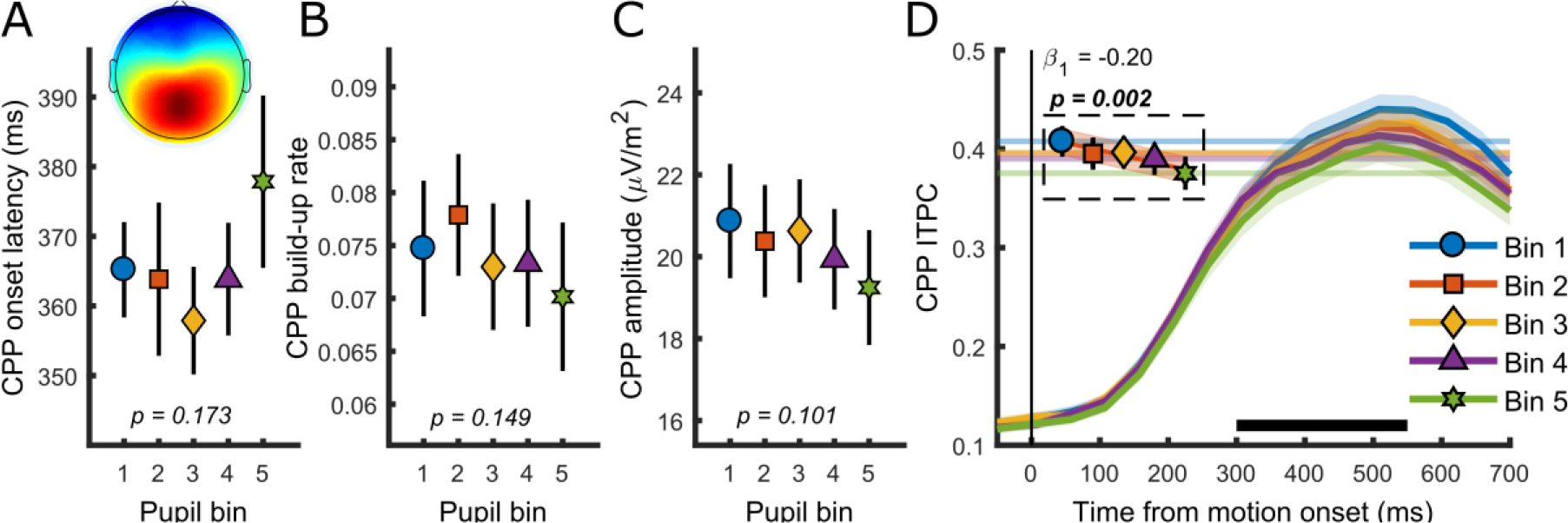
Relationship between baseline pupil diameter and the CPP. (A) CPP onset latency, (B) build-up rate, (C) amplitude and (D) ITPC per pupil bin over time for frequencies below 4 Hz. The black bar indicates the time window used for further analysis. Horizontal lines and symbols indicate the averaged ITPC in the time-frequency window indicated by the white box in Figure 2 panel C. Lines and shading indicate significant fits to the data. Linear fits are displayed when a first-order polynomial fit was superior to a constant fit. Error bars and shaded regions denote ±1 SEM. Stats, linear mixed effects model analyses (Statistical analyses).

Lastly, we investigated whether baseline pupil diameter affected our early target selection signal, the N2 (Figure 6C-D). Previous studies have revealed that baseline pupil diameter affected the size and variability of neural responses to visual and auditory stimuli (Reimer et al., 2014; McGinley et al., 2015a). Here we found that baseline pupil diameter was not predictive of the peak latency of the N2c (p = 0.74), but that it did display a monotonic relationship with the N2c amplitude (χ^2^_(1)_ = 14.31, p < 0.001). Trials with larger baseline pupil diameter displayed smaller N2c amplitudes, suggesting that higher arousal has a negative impact on sensory encoding. N2c ITPC did not vary with baseline pupil diameter (p = 0.30), and nor did N2i ITPC (p = 0.26), N2i latency (p = 0.87) or amplitude (p = 0.06). We thus found that, similar to the phasic pupil diameter response, baseline pupil diameter is predictive of specific characteristics of each of the processing stages of perceptual decision-making. Next, we investigated which of these components explained unique variance in task performance across pupil size bins.

**Figure 6.**
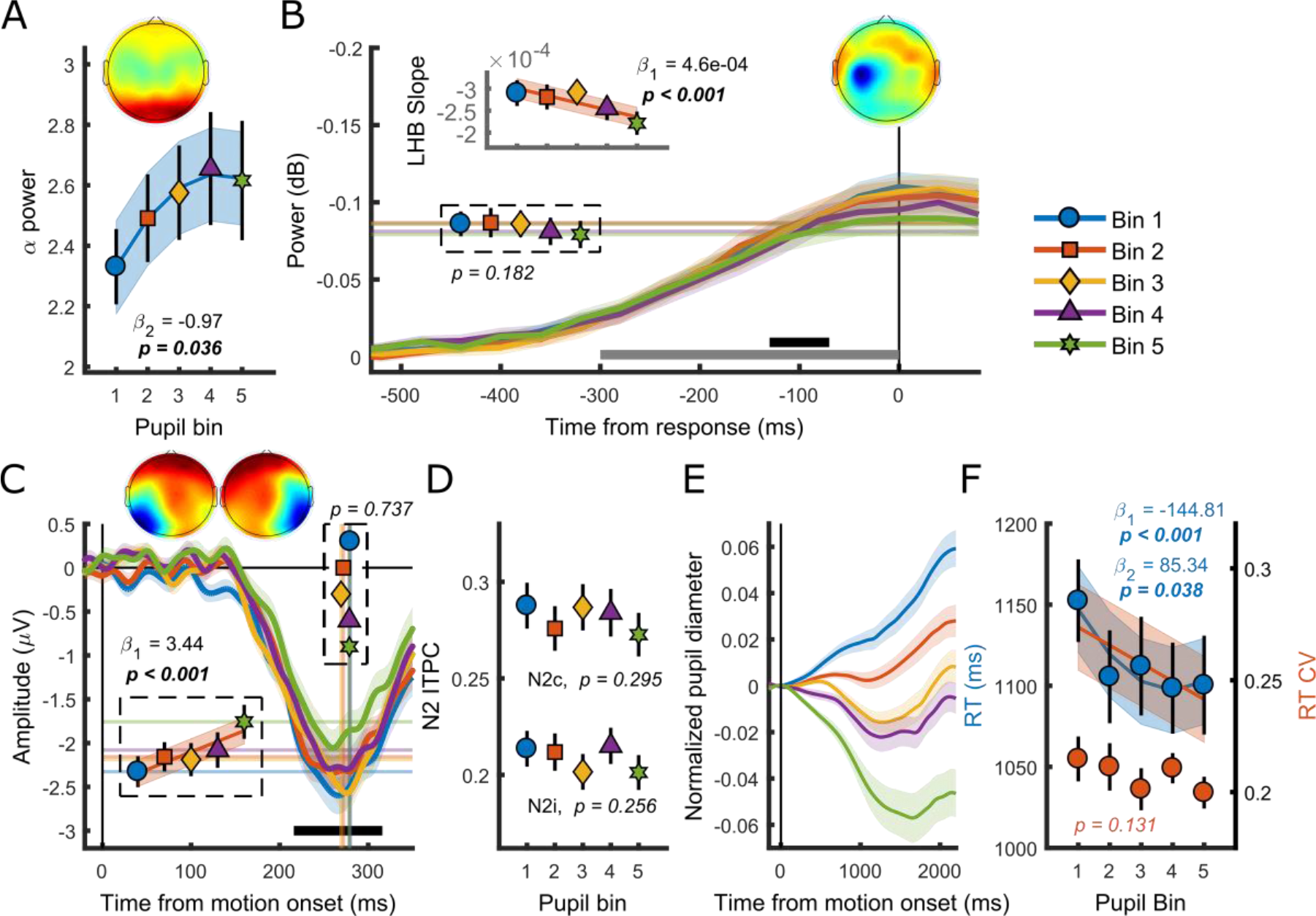
(A) Pre-target α power by baseline pupil diameter. (B) Response-related left hemispheric β power (LHB) per pupil bin. Horizontal lines and markers indicate the average LHB in the time window indicated by the black bar. Inset shows the LHB build-up rate, as determined by fitting a straight line through the LHB in the time window indicated by the grey bar. Note the reverse y-axis direction. (C) The stimulus-locked N2c time-course binned by baseline pupil diameter. Vertical lines and markers show the peak latencies. Horizontal lines and markers show the average N2c amplitude during the time period indicated by the black bar. (D) N2c and N2i ITPC per pupil bin averaged in a time-frequency window based on the grand average (Supplementary figure 2). (E-F) Data from a different dataset using a contrast change detection paradigm where subjects monitored a single central target stimulus (Loughnane et al., 2018). (E) Pupil time-courses for the 5 baseline pupil diameter bins. (F) Relationship between behavioural performance and baseline pupil diameter. Markers indicate mean reaction times (RT, blue, left y-axis) and reaction time coefficient of variation (RTcv, red, right y-axis). Insets show the scalp topography of each neural signal. Lines and shading indicate significant fits to the data. Linear fits are displayed when a first-order polynomial fit was superior to a constant fit, and quadratic fits are displayed where second-order fits were superior to a first-order fit. Error bars and shaded regions denote ±1 SEM. Stats, linear mixed effects model analyses (Statistical analyses).

### Consistency in evidence accumulation mediates the influence of tonic arousal on task performance

We again performed the same hierarchical regression analysis as described above, to see which of the neural signals explained unique variability in task performance associated with tonic arousal. The full results of this analysis are summarised in Supplementary Table 3. Here we discuss the main findings. After the application of a forward/backward model selection algorithm (Venables and Ripley, 2002), N2c amplitude and CPP ITPC were the only parameters that were predictive of RT (Table 1). These variables were forced into one regression model predicting RT, and comparison against a baseline model with baseline pupil diameter as a factor revealed a significant fit (χ^2^_(1)_ = 32.6, p < 0.001) with a marginal (conditional) r^2^ of 4.2% (94.4%). This same hierarchical regression procedure revealed that CPP ITPC was the only EEG component that explained unique variability in RTcv (Table 1). Comparison against a baseline model also led to a significant fit (χ^2^_(1)_ = 26.59, p < 0.001), with a marginal (conditional) r^2^ of 11.7% (43.3%).

Thus, additional to a small effect of N2c amplitude on RT, the consistency of the evidence accumulation process was the only stage of information processing that explained unique within and across-subject variability in task performance associated with changes in baseline pupil diameter.

### During decision-making, baseline pupil diameter does not always predict task performance in a U-shaped manner

Other than a small non-monotonic relationship with pre-target α power, none of the relationships between baseline pupil diameter and the other EEG components was best described by a quadratic polynomial. We therefore asked whether the U-shaped relationship with task performance is a general phenomenon during decision-making. To this end, we again analysed the data from a different dataset using a contrast change detection paradigm where subjects monitored a single central target stimulus (Loughnane et al., 2018). Here, we found a small non-monotonic relationship between baseline pupil diameter and RT (χ^2^_(1)_ = 4.33, p = 0.038), and no relationship with RTcv (p = 0.13) (Figure 6E-F).

Because of the small size of the non-monotonic effect with RT, we repeated this analysis in single-trial, non-binned data, to investigate whether this effect arose from the binning procedure, or time-on-task effects (Supplementary Table 1). This analysis revealed a monotonic relationship between baseline pupil diameter and RT (χ^2^_(1)_ = 8.21, p = 0.004), but no non-monotonic relationship (p = 0.18). Because the non-monotonic relationship was not found using single trial data, we additionally plotted the inverse monotonic relationship between baseline pupil diameter and RT for the binned data (Figure 6F).

It thus seems that on this task, higher levels of central arousal, as opposed to intermediate levels, are associated with improved task performance.

## Discussion

Here we investigated whether behavioural and neural correlates of decision-making varied as a function of baseline or task-evoked pupil diameter. The perceptual decision-making paradigm employed (Figure 1A) allowed us to monitor the relationship between pupil diameter and independent measures of attentional engagement, early target selection, evidence accumulation and motor output. We found that the trial-by-trial variability in both tonic and phasic arousal, as measured by the size of the baseline pupil diameter and pupil response (Figure 1B-D), respectively, were predictive of behavioural performance. This relationship was best described by a second-order, U-shaped, polynomial fit for both RT as well as the variability of RT, RTcv (Figure 1C-E).

We furthermore established that both tonic and phasic arousal were predictive of a subset of EEG signatures, together reflecting discrete aspects of information processing underpinning perceptual decision-making. A hierarchical regression analysis allowed us to determine which of these processing stages exerted an independent influence on behavioural performance associated with central arousal. We found that pre-target α power, indexing baseline attentional engagement, and the build-up rate and consistency of the CPP, reflecting the evidence accumulation process, each explained unique variability in task performance that was due to variability in phasic arousal. Variability in task performance due to variability in tonic arousal, was explained by the amplitude of the target selection signal N2c and the consistency of the CPP.

We thus revealed a direct relationship between both tonic and phasic measures of arousal, and a distinct but overlapping set of EEG signatures of perceptual decision-making.

### Why does the phasic pupil response predict performance in a U-shaped fashion?

Although previous studies have related the size of phasic pupil dilations to behavioural performance (Beatty, 1982b; Kristjansson et al., 2009; de Gee et al., 2017), the association between pupil dilation and speed of detection or cognitive effort is not usually described by a non-linear relationship (but see de Gee et al. (2017) for one account of a non-monotonic relationship with perceptual sensitivity). In this study, we found a strong non-monotonic, U-shaped, relationship between phasic pupil dilations, behavioural performance and EEG signatures during a decision-making task. Here, the largest pupillary constriction and dilation were associated with the poorest behavioural performance, whereas a modest dilation was associated with the best performance.

Arousal determines the way a subject interacts with its environment. Intermediate arousal allows for optimal interaction with the task at hand, whereas suboptimal performance is observed when the subject is either too drowsy or too excitable/distractible (Yerkes and Dodson, 1908; Aston-Jones and Cohen, 2005). At the neuronal level this entails that too little or too much neuromodulatory drive is detrimental for neural signalling and cognition, a phenomenon described for a variety of neuromodulators (Aston-Jones et al., 1999; Aston-Jones and Cohen, 2005; Vijayraghavan et al., 2007; Cano-Colino et al., 2014; Smucny et al., 2016). Although U-shaped relationships between pupil-linked arousal and behavioural performance have previously been found with tonic, rather than phasic measures of central arousal (Aston-Jones and Cohen, 2005; Murphy et al., 2011; McGinley et al., 2015a), the effect of neuromodulatory drive on target neurons after phasic activation could follow a similar U-shaped relationship. Indeed, the classically cited study revealing U-shaped relationships between stimulus intensity and discrimination learning rate investigated whether learning to choose a white over a black passage-way depended on the strength of a “disagreeable electric shock” (Yerkes and Dodson, 1908). Presumably, this shock elicited, amongst others, phasic activation in neuromodulatory arousal centres, which could have influenced the speed with which mice “acquired the habit of avoiding the black-passage way”. Additionally, phasic activation of neuromodulatory systems likely leads to a larger instantaneous increase in neuromodulator availability within or nearby the synaptic cleft than tonic activity (Florin-Lechner et al., 1996; Berridge and Waterhouse, 2003). Phasic arousal could therefore affect target structures and behaviour more strongly and selectively than tonic arousal. Because neuromodulator availability increases transiently upon phasic activation, and these modulators can rapidly be removed from the synaptic cleft (Sarter et al., 2009), target structures could also be less affected by the effects of adaptation for instance, and would thus not display sensitivity decreases to neuromodulators that might be expected during tonic stimulation. Phasic, versus tonic, activity could thus lead to a more local modulator release that supports attentional processes rather than global brain states per se (Thiele and Bellgrove, 2018). Rich computational models that take into account multiple timescales, potential co-transmitters, neuromodulator interactions, and internal behavioural states, as well as input integration from different brain regions (Gjorgjieva et al., 2016), combined with decision-making (O’Connell et al., 2018) could shed light on the neural mechanisms underlying the differential effects of tonic versus phasic neuromodulation on its target structures.

As noted in the Results, we initially hypothesized that the association between low behavioural performance and large pupil responses was due to our use of multiple competing stimuli in which the target location was spatially unpredictable. Kristjansson et al. (2009) compared pupillary responses accompanying slow versus normal performance in a visual vigilance task, focusing on phasic attentional lapses, rather than tonic performance decrements. Subjects monitored three 4-digit timers and were required to indicate as quickly as possible when one of the timers started counting. Slow responses, compared to normal responses, were associated with larger phasic pupil responses. On the other hand, on trials with normal response latencies, the pupil diameter hardly changed. Although pupillary responses on trials with fast RTs were not described, these results do indicate that poor performance can be associated with large pupil dilations on a task that requires monitoring multiple visual stimuli. The required attentional shift could elicit pupil dilations through its relationship with the superior colliculus (Wang et al., 2012; Wang and Munoz, 2015; Joshi et al., 2016; de Gee et al., 2017). On this task, the occurrence of these attentional shifts could be indicated by the amplitude of the N2i. The amplitude of the N2i, ipsilateral to the target and contralateral to the distractor stimulus, was larger for large pupil responses (Figure 4A). This difference was present as early as 70ms after target onset, making it unlikely that it reflects target processing. Rather, large pupil responses, accompanied by larger N2i amplitudes, could indicate that attention was more biased towards one of the stimuli. Trials where the non-attended stimulus turned out to be the target would require a shift in attention, which in turn could be the cause of the delay in response. Indeed, we found an inverse relationship between the N2i amplitude and the size of the pupil response (Figure 4B), suggesting that the need for an attentional shift elicits large pupil responses, which could explain the lower behavioural performance on trials with larger pupil dilations. To see whether attentional shifts could be the sole mechanism by which to explain the U-shaped relationship between pupil response size and task performance, we analysed data from a different experiment in which participants monitored a single stimulus (Figure 4C-E). On this task, although pupil responses did not relate to RTcv, they were predictive of RT in a quadratic manner. The lack of relationship with RTcv on the contrast change detection task implies that variability in RT on the motion detection task could be brought about by shifts in attention, and thus explain the U-shaped relationship between the pupil response and RTcv. However, the quadratic relationship with RT indicates that shifts in attention cannot be the sole cause of the U-shaped relationship between the pupil response and task performance, and that this might be a more general phenomenon during protracted decision-making.

Part of our results can be interpreted in light of the relationship between pupil dilations and the activity in brain areas such as the LC or BF (Aston-Jones and Cohen, 2005; Gilzenrat et al., 2010; Varazzani et al., 2015; Joshi et al., 2016; Reimer et al., 2016; de Gee et al., 2017). Poor performance upon pupil constrictions is in line with studies showing that sensory target detection is suboptimal when a transient LC or BF response is absent (Aston-Jones et al., 1994; Rajkowski et al., 1994; Parikh et al., 2007; Gritton et al., 2016). Additionally, naturally occurring pupillary constrictions are preceded by transient activity decreases in the LC (Joshi et al., 2016), and are associated with increased synchronization of cortical activity, a signature of cortical down states, as well as suboptimal processing of visual stimuli (Reimer et al., 2014). Our results suggest that event-related pupillary constrictions could be associated with similar neural mechanisms.

We additionally found, however, that the largest pupil responses, presumably reflecting the largest phasic modulatory responses, were also associated with lower behavioural performance. Although large pupil responses on trials with low task performance have been reported before (Kristjansson et al., 2009; Murphy et al., 2011; Hong et al., 2014), these results do not seem compatible with an interpretation in which the LC is driving this effect. Direct electrophysiological recordings from the LC have revealed a positive correlation between LC phasic activity and behavioural performance on elementary target detection tasks, without indications that a large phasic LC response leads to worse performance (Aston-Jones et al., 1994, 1997, Rajkowski et al., 1994, 2004). Instead, on trials where discrimination is more difficult and RT latencies are longer, the LC response is delayed (Rajkowski et al., 2004), which would bring about a delay in pupil dilation rather than an immediate, larger response. Although at odds with these studies, Muprhy et al., (2011) previously described a similar relationship, in which trials with large pupil responses were preceded by progressively worse performance which was subsequently followed by better task performance. This finding was interpreted as a compensatory mechanism, driven by cortical performance monitoring brain regions that, via a phasic LC response, possibly reflect a reset of the network (Bouret and Sara, 2005) to reengage participants in the task. Another possible neural mechanism that may lead to the same behavioural outcome is that this effect is driven by cholinergic transients that have been hypothesized to signify a switch from a‘signal-detection down’ to a ‘signal-detection up state’, facilitating target detection (Sarter et al., 2016).

Future research will need to determine which (combination of) brainstem nuclei associated with specific aspects of phasic arousal alterations, or other cognitive functions essential for perceptual decision-making, bring about the effects observed in this study.

### Why does baseline pupil diameter predict performance in a U-shaped fashion?

As predicted by the adaptive gain theory (Aston-Jones and Cohen, 2005), we found optimal performance on trials with intermediate baseline pupil diameter. This effect was however only observed when the pupil diameter data was not high-pass filtered (Supplementary figure 1). This indicates that slow changes (<0.01 Hz) in pupil diameter are driving the effects on task performance. U-shaped relationships with task performance have previously been found during auditory target detection tasks (Murphy et al., 2011; McGinley et al., 2015a), but to the best of our knowledge never during visual decision-making paradigms. Indeed, the effects of pupil-linked arousal can have differential effects on activity across different brain regions (McGinley et al., 2015b). For instance, signal-to-noise ratios of sensory responses in auditory cortex peak at intermediate baseline pupil diameter (McGinley et al., 2015a), whereas in visual areas they are larger for higher baseline pupil diameter (Vinck et al., 2015).

Although we found U-shaped relationships with task performance, in line with previous research (Hong et al., 2014), out of all the investigated EEG components, only pre-target α power displayed a small non-monotonic relationship with baseline pupil diameter. Approximately linear relationships were found with pre-target α power asymmetry, N2c amplitude, LHB amplitude and build-up rate, as well as an inverse relationship with CPP ITPC. Of these, only N2c amplitude and CPP ITPC explained within and across subject variability in task performance (Table 1). It thus seems that the effect of tonic arousal on task performance is mainly driven by an approximately linear relationship with target selection and evidence accumulation consistency. This led us to question whether a U-shaped relationship between tonic arousal and task performance on protracted visual decision-making tasks is a more general phenomenon, or heavily dependent on specific aspects of the behavioural paradigm. The absence of a non-monotonic relationship between baseline pupil diameter and task performance during contrast change detection (Figure 6F) suggests the latter. These differences could be driven by different task demands; on simple tasks performance may benefit from increases in arousal, whereas optimal performance on more difficult discrimination tasks could be found with intermediate arousal (Yerkes and Dodson, 1908; McGinley et al., 2015b). RT was however substantially longer on the task where we did not find a U-shaped relationship (compare Figure 1E & Figure 6F), suggesting that this task was more demanding. Alternatively, the relationship between tonic arousal and task performance could be contingent on attentional demands. On tasks with longer RT that require accumulation of evidence across a longer time-period, greater sustained attention is required, which could benefit from increased arousal and would thus predict an inverse linear relationship between baseline pupil diameter and performance (Figure 6F).

Depending on the behavioural paradigm and task demands, the relationship between central arousal, performance and neural activity may take different forms (McGinley et al., 2015b). Membrane potential recordings from sensory and association areas, as well as direct electrophysiological recordings from neuromodulatory brainstem centres during protracted decision-making, are needed to gain further insight in the exact mechanisms that drive the relationship between cortical state, sensory encoding, evidence accumulation and task performance.

### Recruitment of neuromodulators throughout the decision process

The change in pupil diameter during decision-making (Beatty, 1982a; de Gee et al., 2014, 2017; Lempert et al., 2015; Murphy et al., 2016; Urai et al., 2017) suggests that neuromodulators are recruited throughout the decision-making process, reflecting the sustained ramping activity during evidence accumulation (Cheadle et al., 2014; de Gee et al., 2014, 2017). Our results show that phasic arousal affects several components of the decision variable, the onset, build-up rate and in particular the consistency of the evidence accumulation process. Because variability in CPP ITPC was the main determinant of variability in task performance, our results suggest that phasic arousal affects performance on the task at hand mainly by influencing the variability of the accumulation of sensory evidence.

On trials with very short reaction times, during elementary detection tasks, LC phasic responses are more aligned to the response than target onset (Clayton, 2004), and have therefore been hypothesized to aid the alignment of distributed networks to prepare for motor output (Aston-Jones and Cohen, 2005). During decision-making, however, there may be more time for NA (and other modulators) to influence the decision network (Eckhoff et al., 2009; Nomoto et al., 2010), and could thus influence decisions throughout decision formation (Dayan and Yu, 2006; Cheadle et al., 2014; de Gee et al., 2014, 2017), i.e. during evidence accumulation. Indeed, although the size of the pupil response was predictive of both LHB build-up rate and amplitude, these effects were relatively small and neither EEG component explained unique variance in task performance. Rather, evidence accumulation itself was affected by phasic arousal, which in turn explained variability in task performance. The CPP shares many of the same characteristics of the classic P3 EEG component, suggesting that the P3 reflects the decision formation itself, rather than the neural processes occurring before or after (O’Connell et al., 2012; Kelly and O’Connell, 2013; Twomey et al., 2015). Because of the dense LC innervation of the neural areas thought to be its source, the P3 has been hypothesized to reflect the LC phasic response (Nieuwenhuis et al., 2005). It thus seems likely that the CPP, and therefore evidence accumulation, is also influenced by LC activity. Likewise, it seems plausible that ACh affects attentional processes/evidence accumulation in parietal cortex, and thus also the CPP. Unilateral cholinergic deafferentation of parietal cortex reduces the proportions of neurons that respond to cues, whereas it increases the proportion that respond to distractor stimuli (Broussard et al., 2009). Moreover, whereas in control conditions, the neural population that responded to cues or distractors were largely distinct, after deafferentation these populations overlapped substantially, indicating that cholinergic innervation of parietal cortex is essential for distinguishing cue from distractor and thus for selecting and possibly accumulating the appropriate sensory evidence.

### Variability in task performance due to pupil-linked arousal is best predicted by the consistency in evidence accumulation

During epochs of quiet wakefulness, membrane potential fluctuations of neurons in visual, somatosensory and auditory cortex are closely tracked by baseline pupil diameter (Reimer et al., 2014; McGinley et al., 2015a). These fluctuations in subthreshold membrane potential are characteristic of changing cortical state. Small pupil diameter is characterized by prominent low-frequency (2-10 Hz) and nearly absent high-frequency oscillations (30-80 Hz), whereas larger pupil diameter is characterized by reduced low-frequency, but increased high-frequency oscillations (McGinley et al., 2015a, 2015b). Thus, the average subthreshold membrane potential is most stable during intermediate pupil diameter, when neither low nor high-frequency components predominate. States of lower variability are furthermore characterized by more reliable sensory responses, higher spike rates, increased neural gain and better behavioural performance (Reimer et al., 2014; McGinley et al., 2015a, 2015b). In addition to activity in early sensory areas, there is some evidence that activity in higher-order association areas is also more reliable with intermediate arousal. During auditory target detection, human subjects displayed the least variable RT at intermediate baseline pupil diameter, as well as the highest amplitudes of the P3 component elicited by task-relevant stimuli (Murphy et al., 2011).

Here we found that the consistency of evidence accumulation was the main EEG predictor of variability in task performance associated with both tonic and phasic arousal. For tonic arousal, although CPP ITPC did not follow the same U-shaped relationship as task performance, our findings are largely in line with modelling studies which suggested that higher arousal is specifically predictive of more variability in evidence accumulation (Murphy et al., 2014b). For phasic arousal, higher consistency, and thus less variability, was found for intermediate pupil bins, which also displayed the best behavioural performance. These results suggest that similar neural mechanisms of cortical state described for sensory cortex (Reimer et al., 2014; McGinley et al., 2015b, 2015a; Vinck et al., 2015) might also affect neurons in higher-order association areas (e.g. parietal cortex) and thereby influence evidence accumulation and task performance. Simultaneous pupil diameter and membrane potential recordings in parietal cortex during protracted decision-making are needed to confirm this hypothesis.

### Target selection signal amplitude is modulated by pupil-linked arousal

In the present study, we used a paradigm in which two stimuli were continuously presented and target occurrence was both spatially and temporally unpredictable. Successful target detection thus relied on locating and selecting sensory evidence from multiple sources of information. Loughnane et al. (2016) have shown that these early target selection signals, contralateral to the target stimulus (N2c), modulate sensory evidence accumulation and behavioural performance. Although previous studies have characterised the dependence of the quality of sensory responses on fluctuations in cortical state, as measured by baseline pupil diameter (Reimer et al., 2014; McGinley et al., 2015a; Vinck et al., 2015), to the best of our knowledge, the influence of pupil-linked arousal on target selection signals has not been described before. Here, we showed that early target selection signals are modulated by tonic arousal such that larger baseline pupil diameter was predictive of smaller N2c amplitudes (Figure 6C). Moreover, the amplitude of the N2c also explained unique variability in task performance across pupil bins and subjects (Table 1).

At first glance it seems counterintuitive that target selection amplitudes are decreased, whereas visual encoding in early visual cortex is enhanced on trials with larger baseline pupil diameter (Vinck et al., 2015), or during pupil dilation (Reimer et al., 2014). These differences could be due to differences in the nature of the recordings, as these previous studies used invasive electrophysiology and calcium imaging whereas we used scalp EEG, limiting especially the spatial resolution of our analyses that might be necessary to elucidate these effects (e.g. single neuron orientation tuning). Alternatively, they could constitute differential effects of arousal on visual encoding and target selection. More likely, however, they are due to specific task demands, in particular our use of multiple simultaneously presented competing stimuli. Indeed, there is some evidence that an increase in arousal, as measured by pupil diameter, can increase the ability of a distractor to disrupt performance on a Go/No-Go task in non-human primates (Ebitz et al., 2014). At high arousal levels, performance might thus be negatively affected when the task requires the successful suppression of distracting information, i.e. with higher arousal it is more difficult to focus on the task at hand (Aston-Jones and Cohen, 2005; McGinley et al., 2015b). On the current task, it might thus be more difficult to select and process information from one of the two competing stimuli during states of high arousal, leading to reduced N2c amplitude as well as reduced performance.

In addition to the effects on tonic arousal on the N2c, we found that phasic arousal was predictive of the amplitude of the N2i (Figure 3D). However, because this effect was not restricted to the time period around the peak latency, and present from as early as 70 ms (Figure 4A), it is unlikely to reflect target selection (see above). Rather, it seems plausible that these differences reflect differences in the expected location of target presentation. Thus, our observation that phasic arousal was not predictive of any aspect of target selection is broadly consistent with (de Gee et al., 2017), who found that the pupil response was not predictive of sensory responses.

### Concluding remarks

In this study we investigated the relationship between measures of tonic and phasic pupil-linked arousal and behavioural and EEG measures of perceptual decision-making. We found that trial-to-trial variability in both tonic and phasic arousal accounted for variability in task performance and were predictive of a unique, but overlapping, set of neural metrics of perceptual decision-making. These results confirm our hypothesized relationship between pupil diameter and the electrophysiological correlates of evidence accumulation, providing further support for the notion that the neuromodulators that control central arousal are recruited throughout the decision making process. Moreover, the relationships with task performance were best described by a second-order, U-shaped, polynomial model fit, indicating that during decision-making there are optimal levels of both tonic and phasic activity in the (network of) neuromodulatory centres that control central arousal. Although we found that pupil-linked arousal was predictive of EEG correlates associated with attentional engagement, target selection, evidence accumulation and motor output, the effects of arousal on behavioural performance are mainly mediated through the consistency in evidence accumulation.

## Materials and Methods

### Task procedures

Subjects (n=80) and methods are largely overlapping with the details and procedures described elsewhere (Newman et al., 2017). Here we summarise details necessary to understand this study, and we also describe procedures that differ from the previous study. Participants were seated in a darkened room, 56 cm from the stimulus display (21 inch CRT monitor, 85 Hz 1024 × 768 resolution), asked to perform a continuous bilateral variant (O’Connell et al., 2012; Kelly and O’Connell, 2013) of the random dot motion task (Newsome et al., 1989; Britten et al., 1992). Subjects fixated on a central dot while monitoring two peripheral patches of continuously presented randomly moving dots (Figure 1A). At pseudorandom times, an intermittent period of coherent downward motion (50%) occurred in either the left or the right hemifield. Upon detection of coherent motion, participants responded with a speeded right-handed button press. A total of 288 trials were presented over 16 blocks (18 trials per block).

### Data acquisition and preprocessing

Electroencephalogram (EEG) was recorded from 64 electrodes using an ActiveTwo (Biosemi, 512Hz) system at Trinity College Dublin, Ireland or a BrainAmp DC (Brainproducts, 500Hz) at Monash University, Australia. Data were processed using both custom written scripts and EEGLAB functions (Delorme and Makeig, 2004) in Matlab (MathWorks). Noisy channels were interpolated after which the data were notch filtered between 49-51 Hz, band-pass filtered (0.1-35Hz), and rereferenced to the average reference. Data recorded using the Biosemi system were resampled to 500Hz and combined with the data recorded with the Brainproducts system. Epochs were extracted from −800 to 2800 ms around target onset and baselined with respect to −100 to 0 ms before target onset. To minimize volume conduction and increase spatial specificity, for specific analyses the data were converted to current source density (Kayser and Tenke, 2006). We rejected trials from analyses if the reaction times were <150 or >1700 ms after coherent motion onset, or if either the EEG on any channel exceeded 100 mV, or if the subject broke fixation or blinked (Pupillometry) during the analysis period of the trial, the 500 ms preceding target onset for pre-target α power activity or the interval of 100 ms before target onset to 200 ms after the response.

Pre-target α-band power (8-13 Hz), N2 amplitude and latency, CPP onset and build-up rate and response related β-power amplitude and build-up rate were computed largely in the same way as in Newman et al. (2017). Briefly, α-band power was computed over the 500 ms preceding target onset using temporal spectral evolution (TSE) methods (Thut, 2006), and pooled over two symmetrical parietal regions of interest, using channels O1, O2, PO3, PO4, PO7 and PO8. The N2 components were measured at electrodes P7 and P8, ipsi- and contralateral to the target location (Loughnane et al., 2016; Newman et al., 2017), and the CPP was measured at central electrode Pz. These signals were aggregated to an average waveform for each pupil bin and each participant. We determined the latency of the N2c/N2i as the time point with the most negative amplitude value in the stimulus-locked waveform between 150-400/200-450 ms, while N2c/N2i amplitude was measured as the mean amplitude inside a 100 ms window centered on the stimulus-locked grand average peak (266/340 ms) (Loughnane et al., 2016; Newman et al., 2017).

Onset latency of the CPP was measured by performing running sample point by sample point t-tests against zero across each participant’s stimulus-locked CPP waveforms. CPP onset was defined as the first point at which the amplitude reached significance at the 0.05 level for ≥15 consecutive points. Because we decreased our statistical power by binning the trials into 5 bins (see pupillometry), we did not find an onset for every bin for a subset of subjects (baseline pupil diameter: 13 bins over 11 subjects, pupil response: 16 bins over 12 subjects). Because of our use of linear mixed effect analyses, these subjects could still be included in the analysis, with only the missing values being omitted. Both CPP build-up rate and amplitude were computed using the response-locked waveform of the CSD transformed data to minimize influence from negative-going fronto-central scalp potentials (Kelly and O’Connell, 2013). Build-up rate was defined as the slope of a straight line fitted to this signal in the window from −250 ms to −50 ms before response. CPP amplitude was defined as the mean amplitude within the 100 ms before the response.

Response related left hemisphere β-power (LHB, 20-35 Hz) was measured over the left motor cortex at electrode C3 using short-time Fourier transform (STFT) with a 286 ms window size and 20 ms step size (O’Connell et al., 2012; Newman et al., 2017). LHB amplitude was measured from the response-locked waveform in the window from −130 to −70 ms preceding the response, whereas the LHB build-up rate was defined as the slope of a straight line fitted to this same waveform in the 300 ms before the response.

Inter-trial phase coherence (ITPC) was estimated using single-taper spectral methods from the Chronux toolbox (Bokil et al., 2010) and adapted scripts. We used a 256 sample (512 ms) sliding short time window, with a step size of 25 samples (50 ms). This gave us a half bandwidth (W) of 1.95 Hz: W = (K+1)/2T, with K being the number of data tapers, K=1, and T (s) being the length of the time window. Frequencies were estimated from 0.1 to 35Hz.

### Pupillometry

Eye movements and pupil data were recorded using an SR Research EyeLink eye tracker (Eye-Link version 2.04, SR Research/SMI). Blinks were linearly interpolated from 200 ms before to 200 ms after automatically identified blinks, and the interpolated pupil data was then low-pass filtered (< 6 Hz, second order butterworth). Epochs were extracted from −800 to 2800 ms around coherent motion onset. Trials in which fixation errors or blinks occurred within the analysis period, from 100 ms before target onset to 200 ms after response, were excluded from analysis. Fixation errors were defined as gaze deviations of more than 3°. The pupil diameter was normalized by dividing by the maximum pupil diameter on any trial in the analysis window from 100 ms before target onset to 200 ms after the response for each subject, and baselined on a single trial basis. We computed the baseline pupil diameter by averaging the pupil diameter in the 100 ms before target onset.

A scalar measure of the pupil diameter response was computed by taking the difference between the average pupil diameter in the 400 ms surrounding response and the baseline activity from the same trial. Computing the pupil diameter response over a different size time window surrounding response or by using the linear projection (de Gee et al., 2014; Kloosterman et al., 2015) led to similar results. We used linear regression to remove the trial-by-trial fluctuations in single-trial pupil amplitudes that could be due to baseline pupil diameter, inter-trial interval and target side, all factors that are known to influence either the post target pupil response and/or behavioural response times (Kristjansson et al., 2009; de Gee et al., 2014; Kloosterman et al., 2015; Newman et al., 2017).

Next, we binned our behavioural and EEG data according to either the baseline pupil diameter or the post target pupil response into 5 equally sized bins (mean 49.63 ± SEM 0.81 trials, minimum bin size = 20 trials) (Figure 1B & D). The division into 5 bins allowed us to investigate possible quadratic trends in the data.

### Statistical analyses

We used RStudio (RStudio Team (2016). RStudio: Integrated Development for R. RStudio, Inc., Boston, MA URL http://www.rstudio.com) with the package *lme4* (Bates et al., 2015) to perform a linear mixed effects analysis of the relationship between baseline pupil diameter or the pupil response and behavioural measures and EEG signatures of detection. As fixed effects, we entered pupil bin (see Pupillometry) into the model. As random effects, we had separate intercepts for subjects, accounting for the repeated measurements within each subject. We sequentially tested the fit of a monotonic relationship (first-order polynomial) against a baseline model (zero-order polynomial), and a non-monotonic (second-order polynomial) against the monotonic fit by means of maximum likelihood ratio tests, using orthogonal polynomial contrast attributes. The behavioural or EEG measure *y* was modelled as a linear combination of polynomial basis functions of the pupil bins (*X*):

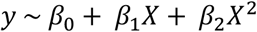
with *β* as the polynomial coefficients. This multilevel approach was preferred over a standard repeated measures analysis of variance (ANOVA), because it allowed us to test for first and second-order polynomial relationships, as well as to account for missing values in the CPP onset estimation. After testing the relationship between behavioural and neural signatures of decision-making and pupillometric measures individually, the neural signals were added sequentially into consecutive regression models predicting RT and RTcv. This model had both a random intercept for each subject, allowing for different baseline-levels of behavioural performance, as well as a random slope of pupil bin, for each subject, which allowed for across-subject variation in the effect of pupil bin on behavioural performance. The hierarchical entry of the predictors allowed us to model the individual differences in behavioural performance, as a function of the EEG signals representing each temporal stage of neural processing. Starting with preparatory signals (α-power), to early target selection signals (N2), to evidence accumulation (CPP), to motor preparation (LHB). The hierarchical addition of the predictors informed us whether each of the EEG signals reflecting successive stages of neural processing improved the fit of the model predicting behavioural data. The signals that explained unique variance were then simultaneously forced into a simplified model predicting RT or RTcv, which made it possible to obtain accurate parameter estimates not contaminated by signals that were shown not to improve model fits. Note that only subjects for which we could determine the CPP onset latency for all bins were included in this hierarchical model. For this final model, all behavioural and neural variables were scaled between 0 and 1 across subjects according to the formula: *y*_*i*_ = (*x*_*i*_ − min *x*_*i*_)/(max *x*_*i*_), where *y*_*i*_ is the scaled variable, *x*_*i*_ is the variable to be scaled. This scaling procedure did not change the relationship of the variable within or across subjects, but scaled all predictor variables to the same range. Again, significance values were obtained by means of maximum likelihood ratio tests.

Data plotted in all figures are the mean and the standard error of the mean (SEM) across subjects. Linear fits are plotted when first-order fits were superior to the zero-order (constant) fit, quadratic fits are plotted when second-order fits were superior to the first-order fit.

### Notes

Raw data (https://figshare.com/s/8d6f461834c47180a444) are open access and available under a Creative Commons Attribution-NonCommercial-ShareAlike 3.0 International Licence. Analysis scripts are freely available on github (https://github.com/jochemvankempen/2018_Monash).

## Acknowledgements

J.K. and A.T. are supported by research funding from the Henry Wellcome Trust (093104). M.A.B. is supported by fellowship and project grant support from the Australian Research Council (ARC; FT130101488; DP150100986; DP180102066). A.T. and M.A.B. are supported by research funding from a strategic research partnership between the Newcastle University and Monash University. M.A.B, R.O.C and A.T are supported by research funding from the Office of Naval Research Global (ONR Global).

## Supplementary information

### The effect of pupil diameter on task performance is not an artefact of the binning procedure

To get an accurate impression of the relationship between pupillary dynamics and task performance, it is best to perform regression analyses on a single trial basis. Unfortunately, many of our behavioural and EEG components of decision-making require the averaging of trials. For instance, RTcv is calculated by dividing the standard deviation by the mean of RT. Likewise, CPP onset latency is computed by performing running sample point by sample point t-tests against zero across each participant’s stimulus-locked CPP waveforms, and cannot be computed on a single-trial basis.

Therefore, we chose to bin our data according to the size of the pupil diameter baseline/response into 5 bins. This however led us to question whether the relationship between pupil diameter and task performance could be dependent on our binning procedure. Therefore, we ran another regression analysis wherein we predicted single trial RT by sequentially adding the linear and quadratic coefficients for baseline pupil diameter (*BPD*) and pupil response (*PR*):

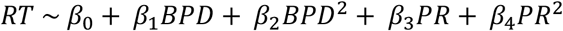
with *β* as the polynomial coefficients. We compared the first model to a random-intercept-only model including subject ID, inter-trial interval, stimulus side, as well as the trial and block number (to control for potential time on task effects), and tested the fit of subsequent models to the previous model fit. This analysis revealed a significant improvement for each step of the sequential analysis, for which the results and parameters estimates are shown in Supplementary Table 1. These analyses confirm that both the size of the baseline pupil diameter and the pupil response are predictive of task performance on a single trial basis. This relationship moreover follows a non-monotonic, quadratic, function.

We repeated this analysis for another dataset in which subjects were required to detect a contrast change in a single centrally presented stimulus (Loughnane et al., 2018). This analysis revealed very similar results, except that the association between baseline pupil diameter and RT was best described by a linear relationship, instead of a second-order polynomial.

**Supplementary Table 1.** Parameter estimates for the single-trial mixed effect model analysis predicting RT using linear and polynomial basis functions of baseline pupil diameter (BPD) and the pupil response (PR). RDM, random dot motion task, CD, contrast change detection task (Loughnane et al., 2018)

|  | Model comparison |  | Parameter estimates |  |  |  |
| --- | --- | --- | --- | --- | --- | --- |
| *RDM* | $\chi^2$ | *p* | $\beta$ | $\beta SE$ | *t* | *p* |
| *BPD* | 5.31 | 0.021 | 0.43 | 0.064 | 6.64 | <0.001 |
| *BPD*<sup>2</sup> | 56.83 | <0.001 | 0.43 | 0.049 | 8.75 | <0.001 |
| *PR* | 65.65 | <0.001 | 0.39 | 0.044 | 8.77 | <0.001 |
| *PR*<sup>2</sup> | 239.67 | <0.001 | 0.72 | 0.046 | 15.53 | <0.001 |
| *CD* |  |  |  |  |  |  |
| *BPD* | 8.21 | 0.004 | -0.14 | 0.027 | -5.14 | <0.001 |
| *BPD*<sup>2</sup> | 1.82 | 0.178 | 0.01 | 0.022 | 0.47 | 0.636 |
| *PR* | 21.70 | <0.001 | -0.12 | 0.023 | -5.16 | <0.001 |
| *PR*<sup>2</sup> | 46.38 | <0.001 | 0.15 | 0.021 | 6.84 | <0.001 |

### Baseline pupil diameter is not predictive of task performance when high-pass filtered

**Supplementary figure 1.**
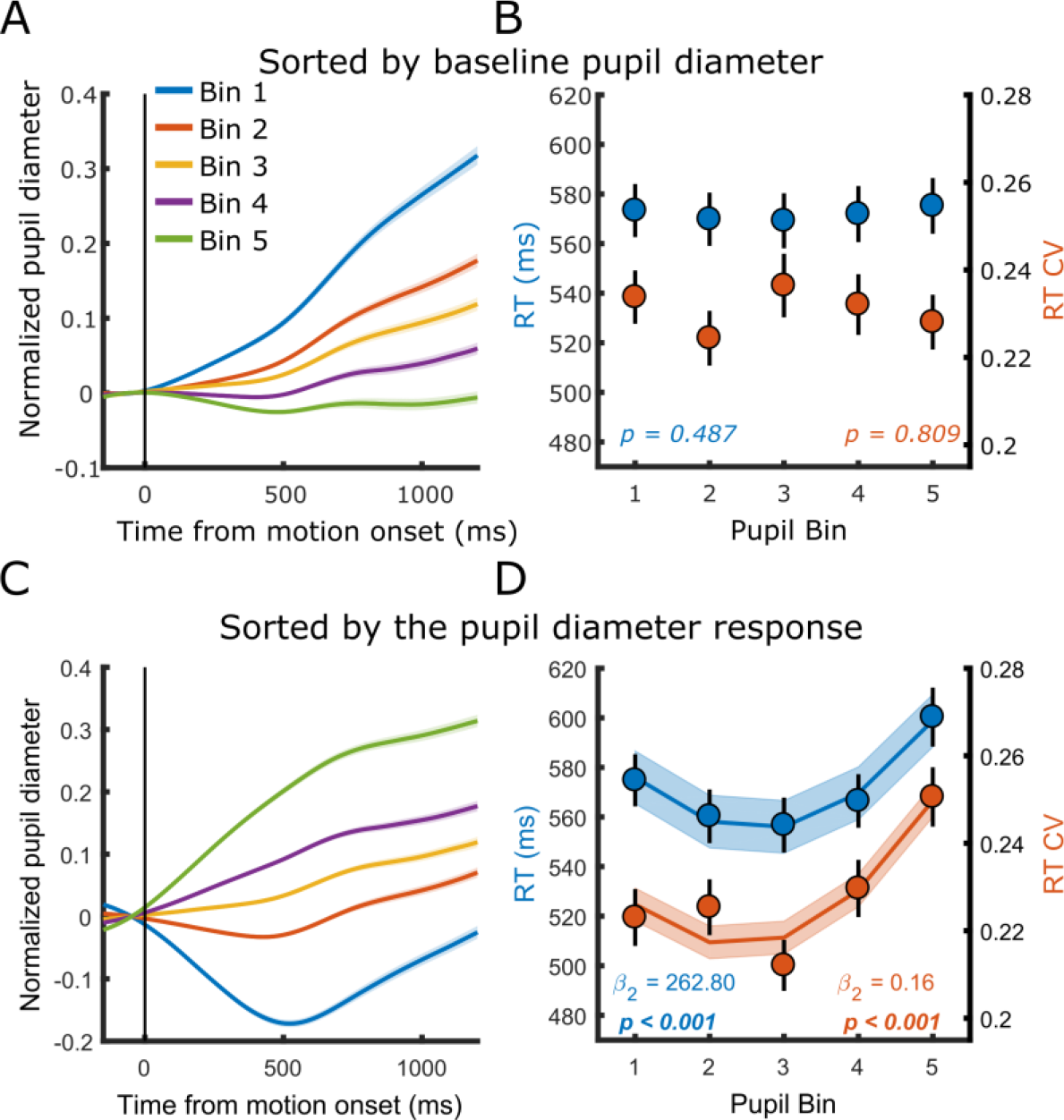
The relationship between baseline pupil diameter and the pupil response with task performance, for band-pass (0.1 - 6 Hz), rather than low-pass (<6Hz) filtered pupil diameter data. (A-B) Pupil diameter time course and task performance sorted by baseline pupil diameter. (A) Pupil time-course for the five bins. (B) Behavioural performance for the five bins. Markers indicate mean reaction times (RT, blue, left y-axis) and reaction time coefficient of variation (RTcv, red, right y-axis). (C-D) Same conventions as A-B, but sorted by the pupil diameter response. Error bars and shaded regions denote ±1 standard error of the mean (SEM). Stats, linear mixed effects model analyses (Statistical analyses).

### N2 ITPC analysis

**Supplementary figure 2.**
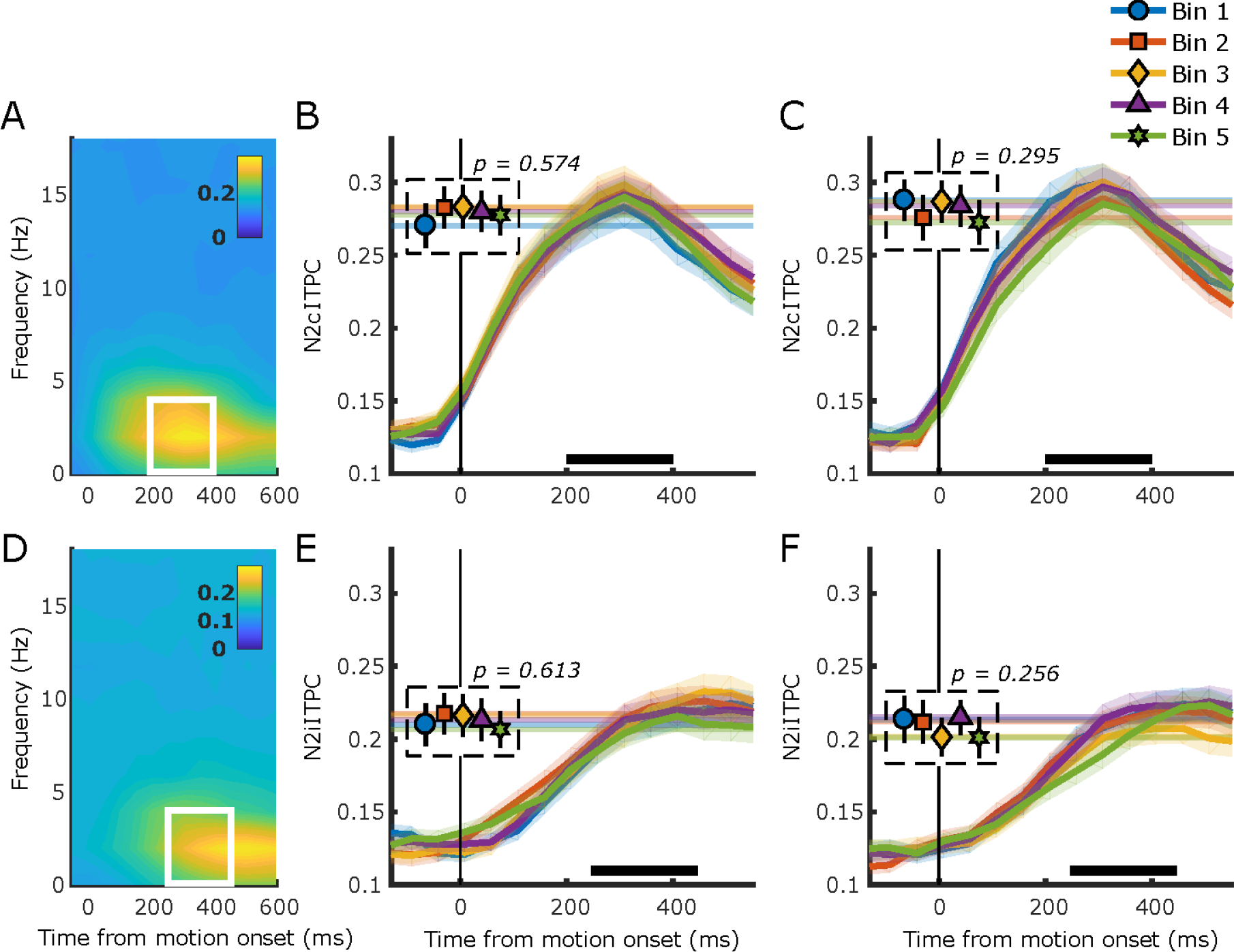
N2 ITPC. (A) Grand average inter-trial phase coherence (ITPC) per time-frequency point for the N2c. White box represents the time-frequency window selected for statistical analyses. (B) N2c ITPC per pupil response bin, (C) N2c ITPC per baseline pupil bin. Horizontal lines indicate average ITPC per pupil bin during the time window indicated by the black bar. (D-F) As in A-C but for N2i. Note that the time window used for the N2i analysis did not cover the peak ITPC activity, but rather focused on the time window in which the N2i amplitude peaked (Figure 3D). Error bars and shaded regions denote ±1 SEM. Stats, linear mixed effects model analyses (Statistical analyses).

### Hierarchical regression analysis predicting variability in task performance associated with phasic and tonic arousal

The tables below show the results from the model comparisons of the hierarchical regression analysis testing which of the signals associated with each of the neural processing stages of perceptual decision-making explained unique variance in task performance associated with phasic (Supplementary Table 2) or tonic arousal (Supplementary Table 3). The neural signals were added to a linear mixed effects model predicting either RT or RTcv in a hierarchical fashion, with their order determined by their temporal order in the decision-making process (Statistical analyses). This allowed us to test whether each successive stage of neural processing would improve the fit of the model to the behavioural data, over and above the fit of the previous stage. To test whether each of the neural signals that significantly improved the model fit indeed explained unique variance in task performance that is not explained by any of the other variables we used an algorithm for forward/backward stepwise model selection (Venables and Ripley, 2002). This procedure could exclude EEG parameters from the final model that for instance are highly correlated to other variables predictive of task performance. The variables that were not eliminated were forced into one linear mixed effects model predicting RT or RTcv, of which the final model parameters are shown in Table 1.

Most of the EEG variables were uncorrelated (r < 0.25). The ones that were correlated were, CPP onset and CPP ITPC (r = 0. 34), CPP amplitude with CPP build-up rate (−0.50) and ITPC (−0.32), and LHB build-up rate and amplitude (−0.33) for baseline pupil diameter, and CPP onset and CPP ITPC (r = 0.43), CPP build-up rate and CPP amplitude (r = −0.59), and LHB build-up rate and amplitude (r = −0.28) for the pupil response.

**Supplementary Table 2.**
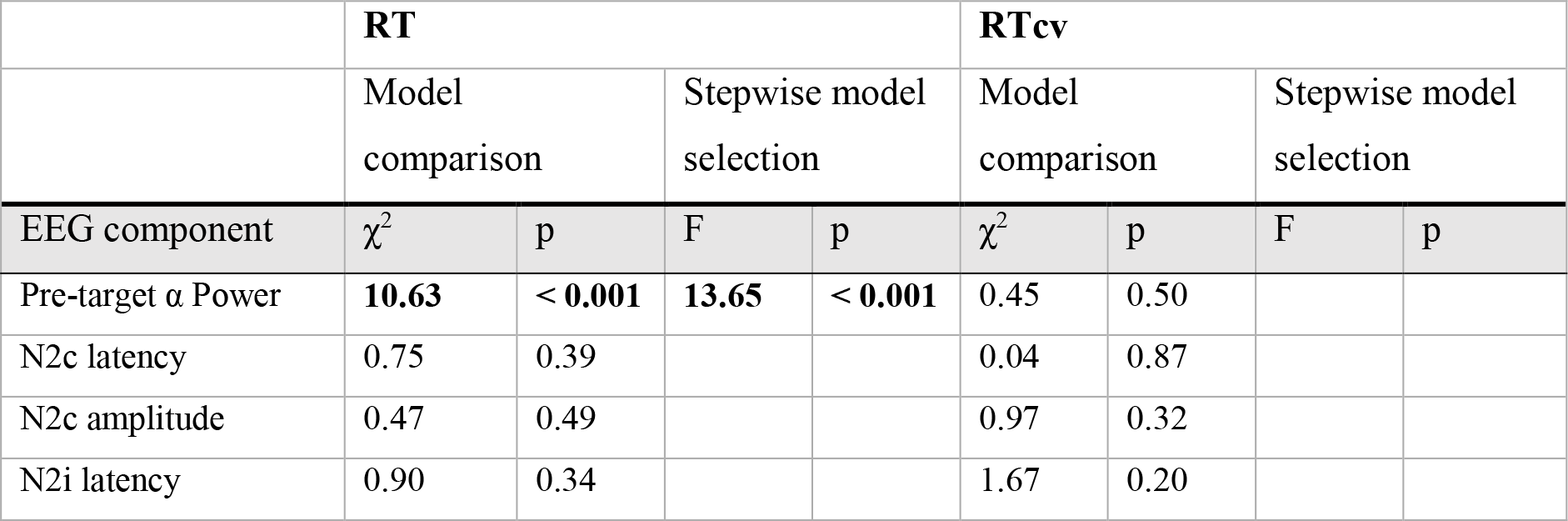
Results from model comparisons of the hierarchical regression analysis predicting variability in task performance due to phasic arousal. Boldface font indicates parameters that significantly improved the model fit compared to the addition of the neural signal associated with the previous neural processing stage. Red text indicates the parameters that were excluded from the final model during the forward/backward stepwise regression (main text). Final model fits revealed a marginal (conditional) r^2^ of 14.6% (93.1%) and 11.1% (46.4%) for RT and RTcv, respectively.

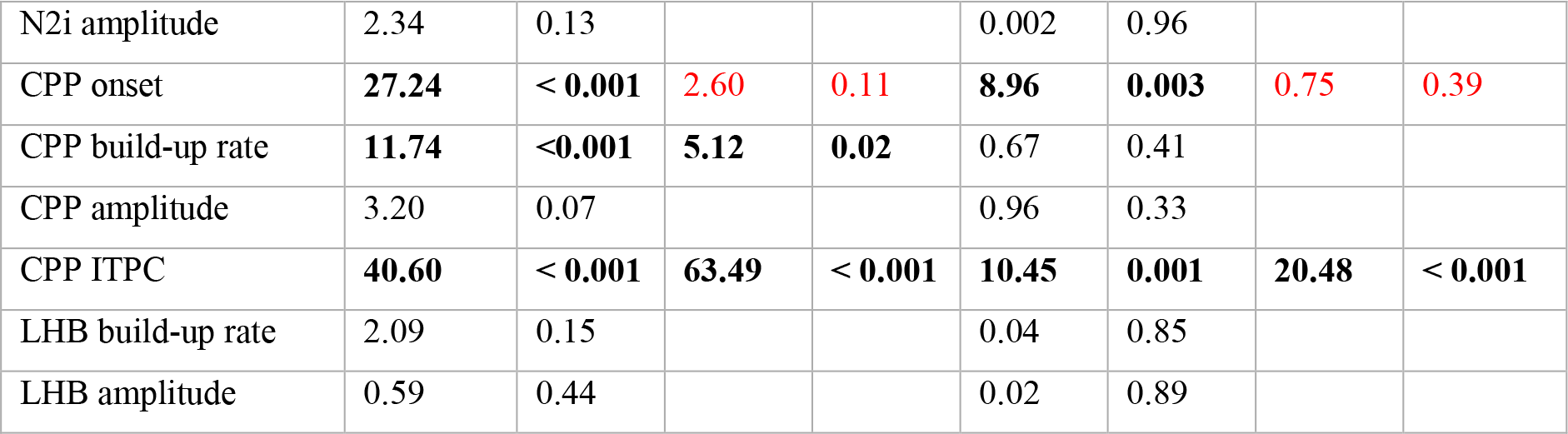

**Supplementary Table 3.**
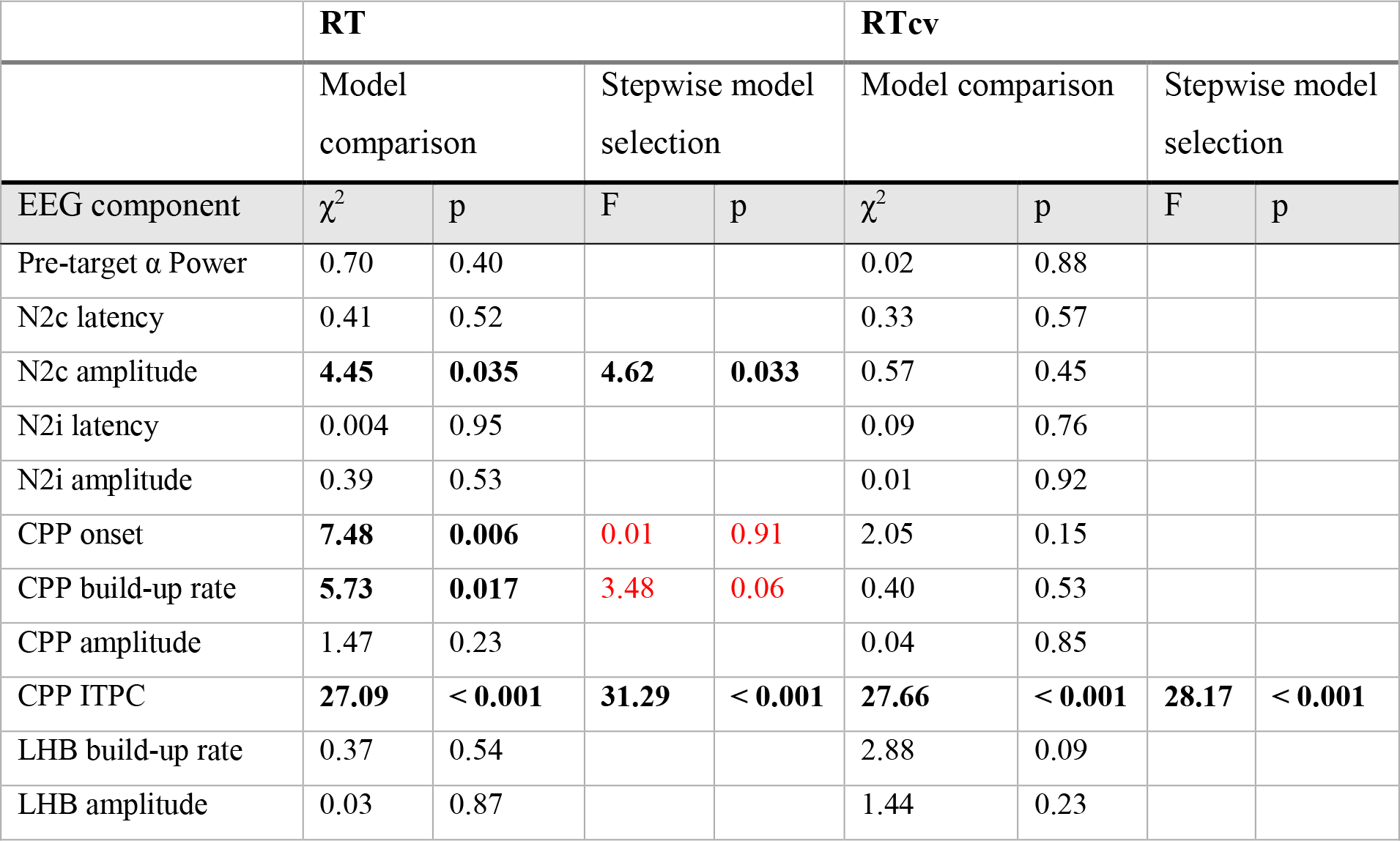
Results from model comparisons of the hierarchical regression analysis predicting variability in task performance due to tonic arousal. Boldface font indicates parameters that significantly improved the model fit compared to the addition of the neural signal associated with the previous neural processing stage. Red text indicates the parameters that were excluded from the final model during the forward/backward stepwise regression (main text). Final model fits revealed a marginal (conditional) r^2^ of 4.2% (94.4%) and 11.7% (43.3%) for RT and RTcv, respectively.

### Robust regression of final parameter estimates predicting variability in task performance

To confirm whether each of the neural signals selected by the hierarchical regression analysis indeed had a significant effect on task performance, we performed a robust regression (Supplementary Table 4) based on 5000 bootstrap replicates to calculate the 95% confidence intervals around the β parameter estimates for the final model fit (Table 1).

**Supplementary Table 4.** Robust regression analysis results. 95% CI for β parameter estimates of the final model fit presented in Table 1.

|  | RT | RTcv |
| --- | --- | --- |
| *Pupil response* |  |  |
| pre-target α-power | 0.083 – 0.280 | |
| CPP build-up rate | -0.181 – -0.012 |  |
| CPP ITPC | -0.287 – -0.172 | -0.373 – -0.148 |
| *Baseline Pupil diameter* |  |  |
| N2c amplitude | 0.002 – 0.107 |  |
| CPP ITPC | -0.241 – -0.114 | -0.416 – -0.196 |

## References

Aston-Jones G, Cohen JD (2005) An integrative theory of locus coeruleus-norepinephrine function: adaptive gain and optimal performance. Annu Rev Neurosci 28:403–450 Available at: http://www.annualreviews.org/doi/10.1146/annurev.neuro.28.061604.135709.

Aston-Jones G, Rajkowski J, Cohen J (1999) Role of locus coeruleus in attention and behavioral flexibility. Biol Psychiatry 46:1309–1320 Available at: http://linkinghub.elsevier.com/retrieve/pii/S0006322399001407.

Aston-Jones G, Rajkowski J, Kubiak P (1997) Conditioned responses of monkey locus coeruleus neurons anticipate acquisition of discriminative behavior in a vigilance task. Neuroscience 80:697–715 Available at: http://linkinghub.elsevier.com/retrieve/pii/S0306452297000602.

Aston-Jones G, Rajkowski J, Kubiak P, Alexinsky T (1994) Locus coeruleus neurons in monkey are selectively activated by attended cues in a vigilance task. J Neurosci 14:4467–4480 Available at: http://www.ncbi.nlm.nih.gov/pubmed/8027789.

Bates D, Mächler M, Bolker B, Walker S (2015) Fitting Linear Mixed-Effects Models Using lme4. J Stat Softw 67 Available at: http://arxiv.org/abs/1406.5823.

Beatty J (1982a) Task-evoked pupillary responses, processing load, and the structure of processing resources. Psychol Bull 91:276–292 Available at: http://doi.apa.org/getdoi.cfm?doi=10.1037/0033-2909.91.2.276.

Beatty J (1982b) Phasic Not Tonic Pupillary Responses Vary With Auditory Vigilance Performance. Psychophysiology 19:167–172 Available at: http://doi.wiley.com/10.1111/j.1469-8986.1982.tb02540.x.

Berridge CW, Waterhouse BD (2003) The locus coeruleus-noradrenergic system: modulation of behavioral state and state-dependent cognitive processes. Brain Res Rev 42:33–84 Available at: http://linkinghub.elsevier.com/retrieve/pii/S0165017303001437.

Bokil H, Andrews P, Kulkarni JE, Mehta S, Mitra PP (2010) Chronux: A platform for analyzing neural signals. J Neurosci Methods 192:146–151 Available at: http://linkinghub.elsevier.com/retrieve/pii/S0165027010003444.

Bouret S, Sara SJ (2005) Network reset: a simplified overarching theory of locus coeruleus noradrenaline function. Trends Neurosci 28:574–582 Available at: http://linkinghub.elsevier.com/retrieve/pii/S0166223605002432.

Britten K, Shadlen M, Newsome W, Movshon J (1992) The analysis of visual motion: a comparison of neuronal and psychophysical performance. J Neurosci 12:4745–4765 Available at: http://eutils.ncbi.nlm.nih.gov/entrez/eutils/elink.fcgi?dbfrom=pubmed&id=1464765&retmode=ref&cmd=prlinks%5Cnpapers3://publication/uuid/E8CEBDDA-F6CF-4392-AEE5-5F9E687F3ED1.

Broussard JI, Karelina K, Sarter M, Givens B (2009) Cholinergic optimization of cue-evoked parietal activity during challenged attentional performance. Eur J Neurosci 29:1711–1722 Available at: http://doi.wiley.com/10.1111/j.1460-9568.2009.06713.x.

Cano-Colino M, Almeida R, Gomez-Cabrero D, Artigas F, Compte A (2014) Serotonin Regulates Performance Nonmonotonically in a Spatial Working Memory Network. Cereb Cortex 24:2449–2463 Available at: https://academic.oup.com/cercor/article-ookup/doi/10.1093/cercor/bht096.

Cheadle S, Wyart V, Tsetsos K, Myers N, de Gardelle V, Herce Castañón S, Summerfield C (2014) Adaptive Gain Control during Human Perceptual Choice. Neuron 81:1429–1441 Available at: http://linkinghub.elsevier.com/retrieve/pii/S0896627314000518.

Clayton EC (2004) Phasic Activation of Monkey Locus Ceruleus Neurons by Simple Decisions in a Forced-Choice Task. J Neurosci 24:9914–9920 Available at: http://www.jneurosci.org/cgi/doi/10.1523/JNEUROSCI.2446-04.2004.

Dayan P, Yu AJ (2006) Phasic norepinephrine: A neural interrupt signal for unexpected events. Netw Comput Neural Syst 17:335–350 Available at: http://www.tandfonline.com/doi/full/10.1080/09548980601004024.

de Gee JW, Colizoli O, Kloosterman NA, Knapen T, Nieuwenhuis S, Donner TH (2017) Dynamic modulation of decision biases by brainstem arousal systems. Elife 6:1–36 Available at: http://elifesciences.org/lookup/doi/10.7554/eLife.23232.

de Gee JW, Knapen T, Donner TH (2014) Decision-related pupil dilation reflects upcoming choice and individual bias. Proc Natl Acad Sci 111:E618–E625 Available at: http://www.pnas.org/lookup/doi/10.1073/pnas.1317557111.

Delorme A, Makeig S (2004) EEGLAB: an open source toolbox for analysis of single-trial EEG dynamics including independent component analysis. J Neurosci Methods 134:9–21 Available at: http://ac.els-cdn.com/S0165027003003479/1-s2.0-S0165027003003479-main.pdf?_tid=336a8184-2684-11e7-8605-00000aab0f02&acdnat=1492773669_497ab65b5f857280a6ed4d13928ea2ca [Accessed April 21, 2017].

Donner TH, Siegel M, Fries P, Engel AK (2009) Buildup of Choice-Predictive Activity in Human Motor Cortex during Perceptual Decision Making. Curr Biol 19:1581–1585 Available at: https://ac.els-cdn.com/S0960982209015437/1-s2.0-S0960982209015437-main.pdf?_tid=089ae988-f9fd-11e7-9ed1-00000aacb362&acdnat=1516025262_164ba055d16590bda0781c671aaacf39 [Accessed January 15, 2018].

Ebitz RB, Pearson JM, Platt ML (2014) Pupil size and social vigilance in rhesus macaques. Front Neurosci 8 Available at: http://journal.frontiersin.org/article/10.3389/fnins.2014.00100/abstract.

Eckhoff P, Wong-Lin KF, Holmes P (2009) Optimality and Robustness of a Biophysical Decision-Making Model under Norepinephrine Modulation. J Neurosci 29:4301–4311 Available at: http://www.jneurosci.org/cgi/doi/10.1523/JNEUROSCI.5024-08.2009.

Eldar E, Cohen JD, Niv Y (2013) The effects of neural gain on attention and learning. Nat Neurosci 16:1146–1153 Available at: http://www.pubmedcentral.nih.gov/articlerender.fcgi?artid=3725201&tool=pmcentrez&rendertype=abstract.

Engel TA, Steinmetz NA, Gieselmann MA, Thiele A, Moore T, Boahen K (2016) Selective modulation of cortical state during spatial attention. Science (80-) 354:1140–1144 Available at: http://www.sciencemag.org/cgi/doi/10.1126/science.aag1420.

Ergenoglu T, Demiralp T, Bayraktaroglu Z, Ergen M, Beydagi H, Uresin Y (2004) Alpha rhythm of the EEG modulates visual detection performance in humans. Cogn Brain Res 20:376–383 Available at: http://linkinghub.elsevier.com/retrieve/pii/S0926641004000941.

Florin-Lechner SM, Druhan JP, Aston-Jones G, Valentino RJ (1996) Enhanced norepinephrine release in prefrontal cortex with burst stimulation of the locus coeruleus. Brain Res 742:89–97 Available at: http://linkinghub.elsevier.com/retrieve/pii/S0006899396009675.

Gilzenrat MS, Nieuwenhuis S, Jepma M, Cohen JD (2010) Pupil diameter tracks changes in control state predicted by the adaptive gain theory of locus coeruleus function. Cogn Affect Behav Neurosci 10:252–269 Available at: http://www.springerlink.com/index/10.3758/CABN.10.2.252.

Gjorgjieva J, Drion G, Marder E (2016) Computational implications of biophysical diversity and multiple timescales in neurons and synapses for circuit performance. Curr Opin Neurobiol 37:44–52 Available at: http://dx.doi.org/10.1016/j.conb.2015.12.008.

Gold JI, Shadlen MN (2007) The Neural Basis of Decision Making. Annu Rev Neurosci 30:535–574 Available at: http://www.ncbi.nlm.nih.gov/pubmed/17600525 [Accessed July 10, 2014].

Gritton HJ, Howe WM, Mallory CS, Hetrick VL, Berke JD, Sarter M (2016) Cortical cholinergic signaling controls the detection of cues. Proc Natl Acad Sci 113:E1089–E1097 Available at: http://www.pnas.org/lookup/doi/10.1073/pnas.1516134113.

Harris KD, Thiele A (2011) Cortical state and attention. Nat Rev Neurosci 12:509–523 Available at: http://www.nature.com/articles/nrn3084.

Hong L, Walz JM, Sajda P (2014) Your Eyes Give You Away: Prestimulus Changes in Pupil Diameter Correlate with Poststimulus Task-Related EEG Dynamics Hamed S Ben, ed. PLoS One 9:e91321 Available at: http://dx.plos.org/10.1371/journal.pone.0091321.

Ignashchenkova A, Dicke PW, Haarmeier T, Thier P (2004) Neuron-specific contribution of the superior colliculus to overt and covert shifts of attention. Nat Neurosci 7:56–64 Available at: http://www.nature.com/articles/nn1169.

Joshi S, Li Y, Kalwani RM, Gold JI (2016) Relationships between Pupil Diameter and Neuronal Activity in the Locus Coeruleus, Colliculi, and Cingulate Cortex. Neuron 89:221–234 Available at: http://dx.doi.org/10.1016/j.neuron.2015.11.028.

Kayser J, Tenke CE (2006) Principal components analysis of Laplacian waveforms as a generic method for identifying ERP generator patterns: I. Evaluation with auditory oddball tasks. Clin Neurophysiol 117:348–368 Available at: http://psychophysiology.cpmc.columbia.edu/pdf/kayser2005b.pdf [Accessed October 19, 2017].

Kelly SP, O’Connell RG (2013) Internal and External Influences on the Rate of Sensory Evidence Accumulation in the Human Brain. J Neurosci 33:19434–19441 Available at: http://www.ncbi.nlm.nih.gov/pubmed/24336710 [Accessed November 18, 2014].

Kloosterman NA, Meindertsma T, van Loon AM, Lamme VAF, Bonneh YS, Donner TH (2015) Pupil size tracks perceptual content and surprise. Eur J Neurosci 41:1068–1078 Available at: http://doi.wiley.com/10.1111/ejn.12859.

Kristjansson SD, Stern JA, Brown TB, Rohrbaugh JW (2009) Detecting phasic lapses in alertness using pupillometric measures. Appl Ergon 40:978–986 Available at: https://ac.els-cdn.com/S0003687009000696/1-s2.0-S0003687009000696-main.pdf?_tid=82ceea64-a927-11e7-b286-00000aacb35e&acdnat=1507137462_930c1e64cea52c7e109b67f751c1e9a9 [Accessed October 4, 2017].

Kustov AA, Lee Robinson D (1996) Shared neural control of attentional shifts and eye movements. Nature 384:74–77 Available at: http://www.nature.com/articles/384074a0.

Latimer KW, Yates JL, Meister MLR, Huk AC, Pillow JW (2015) Single-trial spike trains in parietal cortex reveal discrete steps during decision-making. Science (80-) 349:184–187 Available at: http://www.sciencemag.org/cgi/doi/10.1126/science.aaa4056.

Latimer KW, Yates JL, Meister MLR, Huk AC, Pillow JW (2016) Response to Comment on “Single-trial spike trains in parietal cortex reveal discrete steps during decision-making.” Science (80-) 351:1406–1406 Available at: http://www.sciencemag.org/cgi/doi/10.1126/science.aad3596.

Lee S-H, Dan Y (2012) Neuromodulation of Brain States. Neuron 76:209–222 Available at: http://www.pubmedcentral.nih.gov/articlerender.fcgi?artid=3579548&tool=pmcentrez&rendertype=abstract [Accessed July 11, 2014].

Lempert KM, Chen YL, Fleming SM (2015) Relating Pupil Dilation and Metacognitive Confidence during Auditory Decision-Making Pessiglione M, ed. PLoS One 10:e0126588 Available at: http://journals.plos.org/plosone/article/file?id=10.1371/journal.pone.0126588&type=printable [Accessed August 10, 2017].

Loughnane GM, Newman DP, Bellgrove MA, Lalor EC, Kelly SP, O’Connell RG (2016) Target Selection Signals Influence Perceptual Decisions by Modulating the Onset and Rate of Evidence Accumulation. Curr Biol 26:496–502 Available at: http://dx.doi.org/10.1016/j.cub.2015.12.049.

Loughnane GM, Newman DP, Tamang S, Kelly SP, O’Connell RG (2018) Antagonistic Interactions Between Microsaccades and Evidence Accumulation Processes During Decision Formation. J Neurosci 38:2163–2176 Available at: http://www.jneurosci.org/lookup/doi/10.1523/JNEUROSCI.2340-17.2018.

Lovejoy LP, Krauzlis RJ (2010) Inactivation of primate superior colliculus impairs covert selection of signals for perceptual judgments. Nat Neurosci 13:261–266 Available at: http://www.nature.com/articles/nn.2470.

McGinley MJ, David S V., McCormick DA (2015a) Cortical Membrane Potential Signature of Optimal States for Sensory Signal Detection. Neuron 87:179–192 Available at: http://dx.doi.org/10.1016/j.neuron.2015.05.038.

McGinley MJ, Vinck M, Reimer J, Batista-Brito R, Zagha E, Cadwell CR, Tolias AS, Cardin JA, McCormick DA (2015b) Waking State: Rapid Variations Modulate Neural and Behavioral Responses]. Neuron 87:1143–1161 Available at: http://dx.doi.org/10.1016/j.neuron.2015.09.012.

McPeek RM, Keller EL (2002) Saccade target selection in the superior colliculus during a visual search task. J Neurophysiol 88:2019–2034 Available at: http://jn.physiology.org/content/88/4/2019%5Cnhttp://jn.physiology.org/content/88/4/2019.short%5Cnhttp://jn.physiology.org/content/jn/88/4/2019.full.pdf%5Cnhttp://www.ncbi.nlm.nih.gov/pubmed/12364525.

McPeek RM, Keller EL (2004) Deficits in saccade target selection after inactivation of superior colliculus. Nat Neurosci 7:757–763 Available at: http://www.nature.com/articles/nn1269.

Muller JR, Philiastides MG, Newsome WT (2005) Microstimulation of the superior colliculus focuses attention without moving the eyes. Proc Natl Acad Sci 102:524–529 Available at: http://www.pnas.org/cgi/doi/10.1073/pnas.0408311101.

Murphy PR, Boonstra E, Nieuwenhuis S (2016) Global gain modulation generates time-dependent urgency during perceptual choice in humans. Nat Commun 7:13526 Available at: http://dx.doi.org/10.1038/ncomms13526%5Cnhttp://www.nature.com/doifinder/10.1038/ncomms13526".

Murphy PR, O’Connell RG, O’Sullivan M, Robertson IH, Balsters JH (2014a) Pupil diameter covaries with BOLD activity in human locus coeruleus. Hum Brain Mapp 35:4140–4154 Available at: http://doi.wiley.com/10.1002/hbm.22466.

Murphy PR, Robertson IH, Balsters JH, O’connell RG (2011) Pupillometry and P3 index the locus coeruleus-noradrenergic arousal function in humans. Psychophysiology 48:1532–1543 Available at: http://doi.wiley.com/10.1111/j.1469-8986.2011.01226.x.

Murphy PR, Vandekerckhove J, Nieuwenhuis S (2014b) Pupil-Linked Arousal Determines Variability in Perceptual Decision Making O’Reilly JX, ed. PLoS Comput Biol 10:e1003854 Available at: http://dx.plos.org/10.1371/journal.pcbi.1003854.

Mysore SP, Knudsen EI (2011) The role of a midbrain network in competitive stimulus selection. Curr Opin Neurobiol 21:653–660 Available at: http://linkinghub.elsevier.com/retrieve/pii/S0959438811000924.

Newman DP, Loughnane GM, Kelly SP, O’Connell RG, Bellgrove MA (2017) Visuospatial Asymmetries Arise from Differences in the Onset Time of Perceptual Evidence Accumulation. J Neurosci 37:3378–3385 Available at: http://www.jneurosci.org/content/jneuro/37/12/3378.full.pdf [Accessed April 20, 2017].

Newsome WT, Britten KH, Movshon JA (1989) Neuronal correlates of a perceptual decision. Nature 341:52–54 Available at: http://www.nature.com/doifinder/10.1038/341052a0.

Nieuwenhuis S, Aston-Jones G, Cohen JD (2005) Decision making, the P3, and the locus coeruleus--norepinephrine system. Psychol Bull 131:510–532 Available at: http://dare.ubvu.vu.nl/bitstream/handle/1871/16998/Nieuwenhuis_PsychologicalBulletin_131(4)_2005_u.pdf?sequence=2 [Accessed December 20, 2017].

Nomoto K, Schultz W, Watanabe T, Sakagami M (2010) Temporally Extended Dopamine Responses to Perceptually Demanding Reward-Predictive Stimuli. J Neurosci 30:10692–10702 Available at: http://www.jneurosci.org/cgi/doi/10.1523/JNEUROSCI.4828-09.2010.

O’Connell RG, Dockree PM, Kelly SP (2012) A supramodal accumulation-to-bound signal that determines perceptual decisions in humans. Nat Neurosci 15:1729–1735 Available at: http://www.ncbi.nlm.nih.gov/pubmed/23103963 [Accessed October 8, 2014].

O’Connell RG, Dockree PM, Robertson IH, Bellgrove MA, Foxe JJ, Kelly SP (2009) Uncovering the Neural Signature of Lapsing Attention: Electrophysiological Signals Predict Errors up to 20 s before They Occur. J Neurosci 29:8604–8611 Available at: http://www.jneurosci.org/cgi/doi/10.1523/JNEUROSCI.5967-08.2009.

O’Connell RG, Shadlen MN, Wong-Lin K, Kelly SP (2018) Bridging Neural and Computational Viewpoints on Perceptual Decision-Making. Trends Neurosci xx:1–15 Available at: https://doi.org/10.1016/j.tins.2018.06.005.

Parikh V, Kozak R, Martinez V, Sarter M (2007) Prefrontal Acetylcholine Release Controls Cue Detection on Multiple Timescales. Neuron 56:141–154 Available at: http://ac.els-cdn.com/S0896627307006745/1-s2.0-S0896627307006745-main.pdf?_tid=c3c68606-4baa-11e7-b903-00000aab0f6b&acdnat=1496858425_98631c2399f8a1cfe667358952b60577 [Accessed June 7, 2017].

Parikh V, Sarter M (2008) Cholinergic Mediation of Attention. Ann N Y Acad Sci 1129:225–235 Available at: http://doi.wiley.com/10.1196/annals.1417.021.

Rajkowski J, Kubiak P, Aston-Jones G (1994) Locus coeruleus activity in monkey: Phasic and tonic changes are associated with altered vigilance. In: Brain Research Bulletin, pp 607–616.

Rajkowski J, Majczynski H, Clayton E, Aston-Jones G (2004) Activation of Monkey Locus Coeruleus Neurons Varies With Difficulty and Performance in a Target Detection Task. J Neurophysiol 92:361–371 Available at: http://www.physiology.org/doi/10.1152/jn.00673.2003.

Reimer J, Froudarakis E, Cadwell CR, Yatsenko D, Denfield GH, Tolias AS (2014) Pupil Fluctuations Track Fast Switching of Cortical States during Quiet Wakefulness. Neuron 84:355–362 Available at: http://dx.doi.org/10.1016/j.neuron.2014.09.033 [Accessed October 3, 2017].

Reimer J, McGinley MJ, Liu Y, Rodenkirch C, Wang Q, McCormick DA, Tolias AS (2016) Pupil fluctuations track rapid changes in adrenergic and cholinergic activity in cortex. Nat Commun 7:13289 Available at: http://dx.doi.org/10.1038/ncomms13289.

Sarter M, Lustig C, Berry AS, Gritton H, Howe WM, Parikh V (2016) What do phasic cholinergic signals do? Neurobiol Learn Mem 130:135–141 Available at: http://ac.els-cdn.com/S1074742716000459/1-s2.0-S1074742716000459-main.pdf?_tid=ab1f7ae4-918d-11e7-b172-00000aacb361&acdnat=1504542510_7412be7fc0f08173a5b4462c2ae81331 [Accessed September 4, 2017].

Sarter M, Parikh V, Howe WM (2009) Phasic acetylcholine release and the volume transmission hypothesis: time to move on. Nat Rev Neurosci 10:383–390 Available at: http://www.nature.com/articles/nrn2635.

Shadlen MN, Kiani R, Newsome WT, Gold JI, Wolpert DM, Zylberberg A, Ditterich J, de Lafuente V, Yang T, Roitman J (2016) Comment on “Single-trial spike trains in parietal cortex reveal discrete steps during decision-making.” Science (80-) 351:1406–1406 Available at: http://www.sciencemag.org/cgi/doi/10.1126/science.aaa4056.

Smucny J, Olincy A, Rojas DC, Tregellas JR (2016) Neuronal effects of nicotine during auditory selective attention in schizophrenia. Hum Brain Mapp 37:410–421 Available at: http://doi.wiley.com/10.1002/hbm.23040.

Thiele A, Bellgrove MA (2018) Neuromodulation of Attention. Neuron 97:769–785 Available at: https://doi.org/10.1016/j.neuron.2018.01.008 [Accessed February 27, 2018].

Thut G (2006) -Band Electroencephalographic Activity over Occipital Cortex Indexes Visuospatial Attention Bias and Predicts Visual Target Detection. J Neurosci 26:9494–9502 Available at: http://www.ncbi.nlm.nih.gov/pubmed/16971533 [Accessed October 7, 2014].

Twomey DM, Murphy PR, Kelly SP, O’Connell RG (2015) The classic P300 encodes a build-to-threshold decision variable. Eur J Neurosci 42:1636–1643 Available at: http://doi.wiley.com/10.1111/ejn.12936.

Urai AE, Braun A, Donner TH (2017) Pupil-linked arousal is driven by decision uncertainty and alters serial choice bias. Nat Commun 8:14637 Available at: http://www.nature.com/doifinder/10.1038/ncomms14637.

van Dijk H, Schoffelen J-M, Oostenveld R, Jensen O (2008) Prestimulus Oscillatory Activity in the Alpha Band Predicts Visual Discrimination Ability. J Neurosci 28:1816–1823 Available at: http://www.jneurosci.org/cgi/doi/10.1523/JNEUROSCI.1853-07.2008.

Varazzani C, San-Galli A, Gilardeau S, Bouret S (2015) Noradrenaline and Dopamine Neurons in the Reward/Effort Trade-Off: A Direct Electrophysiological Comparison in Behaving Monkeys. J Neurosci 35:7866–7877 Available at: http://www.jneurosci.org/content/jneuro/35/20/7866.full.pdf [Accessed January 3, 2018].

Venables WN, Ripley BD (2002) Modern Applied Statistics with S, 4th ed. New York: Springer.

Vijayraghavan S, Wang M, Birnbaum SG, Williams G V, Arnsten AFT (2007) Inverted-U dopamine D1 receptor actions on prefrontal neurons engaged in working memory. Nat Neurosci 10:376–384 Available at: http://www.ncbi.nlm.nih.gov/pubmed/17277774 [Accessed November 24, 2014].

Vinck M, Batista-Brito R, Knoblich U, Cardin JA (2015) Arousal and Locomotion Make Distinct Contributions to Cortical Activity Patterns and Visual Encoding. Neuron 86:740–754 Available at: http://dx.doi.org/10.1016/j.neuron.2015.03.028 [Accessed October 2, 2017].

Wang C-A, Boehnke SE, White BJ, Munoz DP (2012) Microstimulation of the Monkey Superior Colliculus Induces Pupil Dilation Without Evoking Saccades. J Neurosci 32:3629–3636 Available at: http://www.jneurosci.org/cgi/doi/10.1523/JNEUROSCI.5512-11.2012.

Wang C-A, Munoz DP (2015) A circuit for pupil orienting responses: implications for cognitive modulation of pupil size. Curr Opin Neurobiol 33:134–140 Available at: http://dx.doi.org/10.1016/j.conb.2015.03.018.

Yerkes RM, Dodson JD (1908) The relation of strength of stimulus to rapidity of habit-formation. J Comp Neurol Psychol 18:459–482 Available at: http://doi.wiley.com/10.1002/cne.920180503 [Accessed January 9, 2018].

